# A novel imprinting cluster at the porcine *CRSP* complex locus defines a species-specific imprinted domain

**DOI:** 10.64898/2026.01.09.698515

**Authors:** Jinsoo Ahn, In-Sul Hwang, Mi-Ryung Park, In-Cheol Cho, Seongsoo Hwang, Kichoon Lee

## Abstract

**Background:** Genomic imprinting is an epigenetic phenomenon that results in parent-of-origin-specific gene expression and has been extensively characterized in mice and humans. However, in pigs, imprinting has been investigated primarily through analyses of orthologs of known imprinted genes in mice and humans. The objective of this study was to examine DNA methylation status and gene expression at a porcine locus containing newly identified imprinted calcitonin receptor-stimulating peptide (CRSP)-encoding genes, to compare orthologous loci in mice and humans, and to investigate a potential underlying mechanism.

**Results:** Analyses of differentially methylated regions (DMRs) between porcine parthenogenetic embryos and biparental controls revealed multiple parental DMRs at a locus we term the *CRSP* complex locus, which harbors CRSP-encoding genes that likely arose through gene duplication. In contrast, orthologous genomic intervals in mice and humans exhibited unmethylated promoters and lacked evidence of imprinting. Consistently, CRSP-encoding genes in pigs showed parent-of-origin-specific monoallelic expression, whereas genes within the orthologous locus in mice and humans were biallelically expressed. Further analysis indicated that porcine *CRSP* promoters are embedded within oocyte-expressed alternative transcripts and co-occurred with DNA methylation, suggesting a transcription-dependent imprinting mechanism.

**Conclusions:** Our comparative analyses identified CRSP-encoding genes at the porcine *CRSP* complex locus as novel imprinted genes, indicating species-specific evolution of this imprinted domain. The results further suggest that lineage-specific gene duplication may have contributed to the emergence of imprinting at this locus.

## Background

Genomic imprinting is an epigenetic phenomenon that leads to parent-of-origin-specific (paternal or maternal) gene expression and plays a crucial role in growth-related traits during mammalian development and growth [1, 2]. The parental conflict theory proposes that paternally expressed genes tend to enhance growth, whereas maternally expressed genes restrict resource allocation [3–5]. As an inherited imprinting mark, DNA methylation is essential for silencing either the maternal or paternal allele during development [6, 7]. The establishment of differentially methylated regions (DMRs) between maternal and paternal alleles in the gametes, and their maintenance after fertilization in the zygote and early embryos, give rise to germline DMRs (gDMRs), which are regarded as primary imprints. These epigenetic marks can function as imprinting control regions (ICRs) that regulate parental allele-specific gene expression of nearby or clustered genes within genomic domains [2]. As these ICRs govern the expression of growth-relevant genes, imprinted genes and their associated regulatory variants serve as valuable genetic markers for selecting livestock with improved growth traits [8]. Nevertheless, despite extensive imprinting research documenting approximately 200 imprinted genes in mice and humans [9, 10], only a relatively small number, around 40, have been cataloged in pigs [10–12]. These observations highlight the critical need to identify additional DMRs/ICRs and imprinted genes that contribute to pig growth and development. Given this need, genomic regions with important physiological roles and lineage-specific gene architecture offer targets for investigating imprinting status.

A human locus on chromosome 11p contains the calcitonin-related polypeptide alpha (*CALCA*) and beta (*CALCB*) genes, which encode the two isoforms of calcitonin gene-related peptide (CGRP), a major target of recently developed migraine-specific therapeutics, including CGRP-related monoclonal antibodies [13–15]. The *CALCA* gene encodes α-CGRP as well as the hormone calcitonin (CT), whereas *CALCB* encodes β-CGRP only, a paralog of α-CGRP. CT released from thyroid C-cells lowers elevated blood calcium levels by inhibiting bone resorption, whereas CGRP functions as a neuropeptide that mediates neural signaling and vasodilation [16, 17]. The arrangement of *Calca* and *Clacb* genes in the corresponding locus in mice is conserved. In contrast, in the orthologous locus in pigs, the calcitonin receptor-stimulating peptides CRSP1, CRSP2, and CRSP3 have been identified as members of the CT/CGRP/CRSP peptide family [18–20]. Although CRSPs show high amino acid sequence similarity with CGRP [21], they exhibit distinct biological properties and have not been identified in human, non-human primates, or rodents [21].

Based on the current pig gene annotation (Sscrofa11.1), CRSP1 is encoded by *CALCB_1*, CRSP2 by *CRSP-2*, and CRSP3 by *CRSP3*, with an additional CRSP1-like peptide encoded by *LOC110259565*. These CRSP-encoding genes (*CALCB_1*, *CRSP-2*, *CRSP3*, and *LOC110259565*) are located within a genomic region that we refer to as the *CRSP* complex locus, situated between *INSC*–*CALCB* and *CYP2R1* in pigs, and notably, the orthologous *CALCA* gene is absent from this region. Functionally, CRSP1 stimulates the porcine calcitonin receptor (CT-R) but not the CGRP receptor (CGRP-R), despite sequence similarity between CRSP1 and CGRP, whereas CRSP2 and CRSP3 stimulate neither CT-R nor CGRP-R [18, 19, 21]. CRSP1 binds CT-R with high affinity and stimulates cAMP production with a potency more than 100-fold greater than CT, which led to its designation as calcitonin receptor-stimulating peptide [18]. Although CT-R is expressed in the central nervous system (CNS) as well as in peripheral tissues, expression of CT itself is absent in the CNS [18]. Because CRSP1 is highly expressed in the CNS, it was hypothesized to function as a cognate ligand for CT-R in the CNS [18]. Despite the evolutionary divergence and specialized biological functions of the CRSP-encoding genes in pigs, whether these genes exhibit parent-of-origin-specific expression remains unknown. To date, no comparative analysis has systematically examined the imprinting status of the genes encoding the CT/CGRP/CRSP peptide family across humans, mice, and pigs.

Here, we aimed to investigate differential methylation within the putative promoter regions of the *CRSP* complex locus between bimaternal parthenogenetic and normal biparental pig embryos using whole-genome bisulfite sequencing (WGBS), and to identify parental DMRs. We detected parental DMRs within the locus spanning the putative promoters of the CRSP-encoding genes. Comparative analysis revealed that, unlike in humans and mice, the pig locus exhibits partial methylation and that the CRSP-encoding genes display parent-of-origin-specific expression, consistent with genomic imprinting. Notably, phylogenetic analysis supported gene duplication within the locus, and the presence of long terminal repeats and oocyte-specific transcription provides mechanistic insights into imprinting establishment. Taken together, these findings highlight the *CRSP* complex locus as a novel imprinted locus in pigs, characterized by multiple parental DMRs, corresponding imprinted gene expression, associated regulatory elements, and an underlying imprinting mechanism.

## Results

### Analysis of parental allele-specific methylation reveals four DMRs in the porcine *CRSP* complex locus

To investigate maternal and paternal DNA methylation in porcine embryos, we generated parthenogenetically activated (PA) embryos and compared them with normal control (CN) embryos. PA embryos inherit bi-maternal alleles, whereas CN embryos inherit bi-parental alleles (maternal and paternal), allowing for allelic comparison. Subsequently, we identified a series of methylation differences between PA and CN embryos on pig chromosome 2 (chr2) in a region located proximal to the centromere (Fig. 1A). Based on the current Sscrofa11.1 gene annotation (GCF_000003025.6; gene transfer format [GTF] file), this region corresponds to a locus located between the *INSC*–*CALCB* and *CYP2R1* genes, which we termed the *CRSP* complex locus. This locus (chr2:44.13‒44.37 Mb) contains four calcitonin receptor-stimulating peptide genes (*CRSP*s): (i) *CRSP3*, (ii) *CRSP-2* (also referred to as *CRSP2* in the literature [20]), (iii) a *CALCB*-like gene, annotated as *CALCB_1* in the GTF file and originally reported as *CRSP* [18] or *CRSP1* [20], and (iv) *LOC110259565*, a predicted calcitonin receptor-stimulating peptide 1-like (*CRSP1-like*) (Fig. 1A). The current annotation does not include *CALCA* (encoding CT/α-CGRP). The locus is flanked distally by *INSC* and *CALCB* (encoding β-CGRP) and proximally by *LOC110259296* (an uncharacterized ncRNA gene) and *CYP2R1*.

**Fig. 1.**
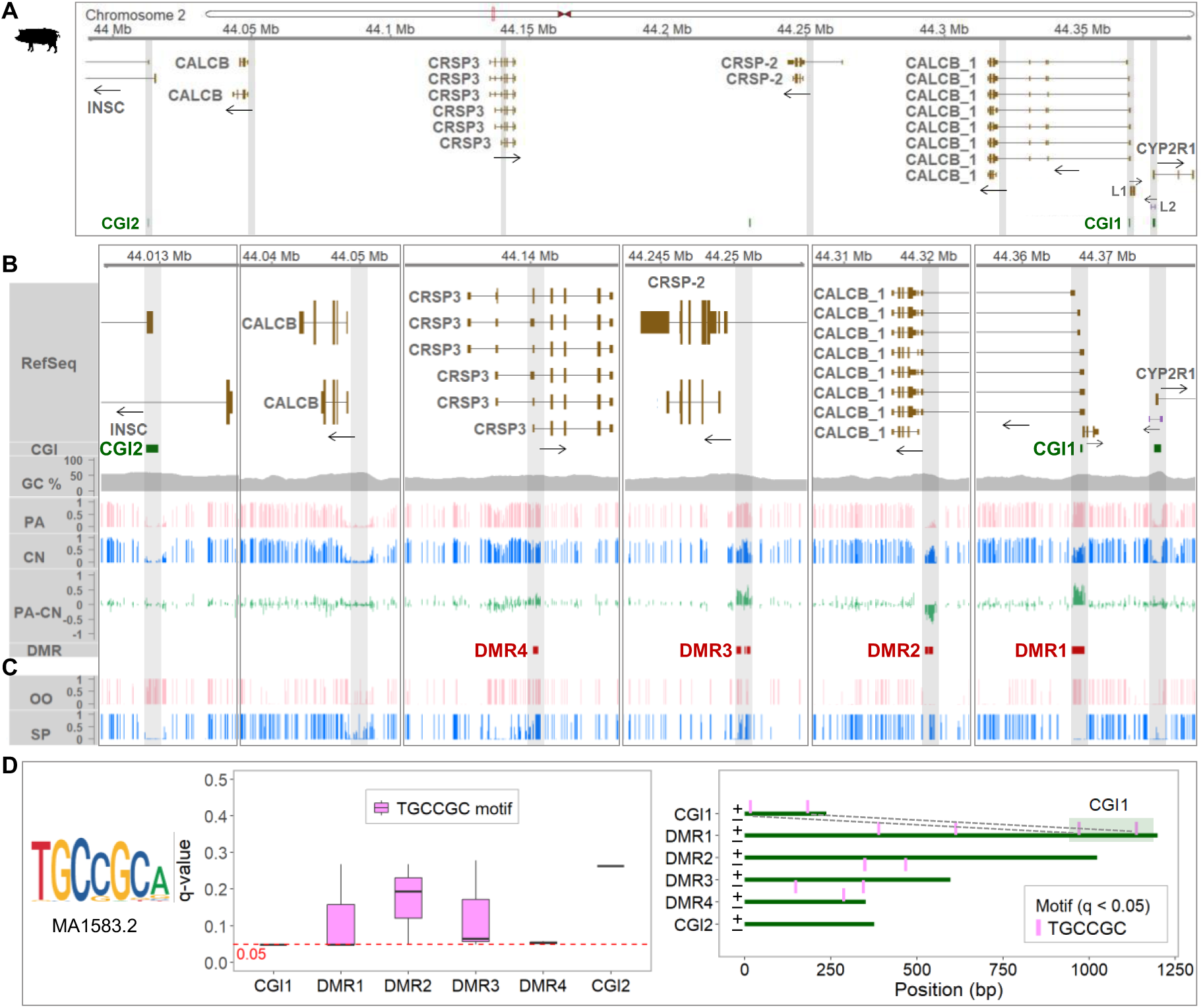
Overview of the porcine *CRSP* complex locus, parental DNA methylation patterns, and associated motif occurrence. **(A)** NCBI RefSeq gene models are shown for the genomic interval chr2:44,000,000‒44,388,000 (Sscrofa11.1), located between *INSC* and *CYP2R1* and encompassing the *CRSP* complex locus (chr2:44.13‒44.37 Mb). Transcriptional directions are indicated by arrows. Upstream of *CALCB_1*, three genes are present in the following order: *LOC110259565* (*CRSP1-like*, L1 in the figure), *LOC110259296* (uncharacterized ncRNA gene, L2 in the figure), and *CYP2R1*. **(B)** CpG methylation profiles for the putative promoter regions highlighted in (A). Mean methylation levels for parthenogenetically activated (PA) and control (CN) embryos, derived from triplicate samples, and their differences (PA – CN) are shown. Differentially methylated regions (DMRs) were identified using metilene (FDR < 0.05). CGI, CpG island; CG%, GC content. **(C)** Germline CpG methylation profiles for porcine oocyte (OO) and sperm (SP) (GEO: GSE143850) are shown for the same genomic interval. **(D)** Analysis of the consensus ZFP57 binding motif (TGCCGC; MA1583.2 from the JASPAR database) in CGIs and DMRs using FIMO. Locations of significantly matched sites (q-value < 0.05) are shown for the + and – strands.

WGBS analysis identified four differentially methylated regions (DMRs; FDR < 0.05), and among them, DMR1, DMR3, and DMR4 were hypermethylated in PA embryos compared with CN embryos (Fig. 1B; Table S1). This pattern suggests that these are maternally methylated DMRs, as bi-maternal PA embryos tended to exhibit approximately double the methylation levels relative to CN embryos carrying a single maternal allele (reflecting a full-versus-half methylation pattern), consistent with loci subject to maternal methylation, although the increase was less pronounced at DMR4. Conversely, DMR2 was hypermethylated in CN embryos compared with PA embryos (Fig. 1B; Table S1). It suggests that DMR2 is a paternally methylated DMR, as DNA methylation was largely absent or present at very low levels in bi-maternal PA embryos but clearly detectable in CN embryos carrying the paternal allele (reflecting a none-versus-half methylation pattern).

The clustered DMR domain encompassing the putative promoter regions of (i) the *CALCB_1* long-form and *LOC110259565*, (ii) *CALCB_1* short-form, (iii) *CRSP-2* short-form, and (iv) *CRSP3* short-form was clearly evident in the DNA methylation landscape between *INSC* and *CYP2R1* (Fig. S1A; Table S1). In contrast, (i) the *CALCB* gene located immediately upstream of *INSC*, (ii) *INSC*, (iii) *CYP2R1*, and (iv) *LOC110259296* (ncRNA) showed typical unmethylated patterns in their putative promoter regions in both PA and CN embryos (Fig. 1B; Fig. S1A; Table S1). Together, these methylation profiles suggest that the maternally and paternally methylated DMRs are associated with promoter regions (promoter-associated DMRs), in contrast to the typical unmethylated promoters observed in both PA and CN embryos. Additionally, the putative promoter regions of the *CRSP-2* long-form and the *INSC* long-form were hypermethylated, a pattern typically associated with transcriptional repression (Fig. 1B; Fig. S1A).

Collectively, these results from the parthenogenetic model highlight a previously unreported DMR domain within the porcine *CRSP* complex locus, which contains a cluster of paralogous calcitonin receptor-stimulating peptide genes, consisting of maternally methylated DMR1, DMR3, and DMR4 and paternally methylated DMR2, which warrants further in-depth imprinting investigation.

### Gametic differential methylation and motif occurrence support the establishment and maintenance of germline DMRs

We further examined DNA methylation in oocytes and sperm and compared their methylation profiles with those of PA and CN embryos to identify germline DMRs and infer their parental origin. The regions overlapping or flanking the maternally methylated DMRs (DMR1, DMR3, and DMR4) were fully methylated in oocytes but unmethylated or methylated at low levels in sperm (Fig. 1B-C; Fig. S1B-C), indicating that these DMRs correspond to the methylation pattern characteristic of maternal germline DMRs and may reflect the persistence of maternal methylation. In particular, this pattern was prominent at DMR1 and DMR3, while DMR4 was associated with a broader region of maternal methylation extending into an adjacent region toward the first exon of the *CRSP3* middle-form transcript (Fig. 1B-C; Fig. S1B-C). Conversely, the region overlapping the paternally methylated DMR2 was fully methylated in sperm, whereas methylation was deficient in oocytes (Fig. 1B-C; Fig. S1B-C), suggesting that DMR2 likely represents a paternal germline DMR and that this paternal methylation appears to be maintained after fertilization.

Additionally, among three CpG islands in this locus, we identified two that are differentially methylated between oocytes and sperm (oocyte methylation and sperm unmethylation; designated as CGI1 and CGI2), whereas the CpG island at the *CYP2R1* promoter region was unmethylated in both gametes (Fig. 1C). Given the distinct methylation patterns observed at CGI1 and CGI2 in embryos (maternal methylation at CGI1 and loss of methylation at CGI2) (Fig. 1B), we investigated the presence of motifs associated with methylation maintenance through motif analysis. FIMO (Find Individual Motif Occurrences) was employed to identify the consensus ZFP57 binding motif (TGCCGC; MA1583.2 from the JASPAR database), which is implicated in methylation maintenance at imprinting control regions (ICRs) [22]. Within CGI1, two predicted motif occurrences matched the TGCCGC sequence and were statistically significant (q = 0.0478), whereas CGI2 contained a variant (GGCCGC) that was not significant (q = 0.263) (Fig. 1D; Table S2). Moreover, all four analyzed DMRs, where DMR2 and DMR3 were each concatenated for FIMO analysis due to their fragmented distributions, harbored at least one statistically significant ZFP57 motif (q < 0.05): four in DMR1 (including two within CGI1), two in DMR2, two in DMR3, and one in DMR4 (Fig. 1D; Table S2), supporting a potential role for ZFP57 in methylation maintenance associated with the maternally methylated (DMR1, DMR3, and DMR4) and paternally methylated (DMR2) germline DMRs.

Taken together, these findings suggest that methylation is established in porcine gametes and potentially maintained after fertilization at the *CRSP* complex locus, forming a novel imprinted domain characterized by multiple germline DMRs.

### Comparative promoter analysis uncovers conserved unmethylation in mice and humans, whereas pigs show partial methylation at the *CRSP* complex locus

To examine the methylation status of the orthologous locus between *INSC* and *CYP2R1* in other mammals, we analyzed somatic tissues from mice and humans and compared the results with those from pigs. The order of protein-coding genes within the murine *Insc‒Cyp2r1* region on chromosome 7 (cytoband 7F1; chr7:114.35‒114.15 Mb, reverse strand) is *Insc*, *Calcb*, *Calca*, and *Cyp2r1*, with no CRSP-encoding genes present (Fig. 2A). Analysis of WGBS data from adult mouse somatic tissues (GSE42836) revealed that putative promoter regions encompassing the transcription start site (TSS) between the *Insc* and *Cyp2r1* genes exhibited a typical unmethylated or markedly hypomethylated pattern across all analyzed tissues (brain [ectoderm], heart and kidney [mesoderm], and liver and lung [endoderm]), representing all three germ layers, whereas the *Insc* long-form promoter region was exceptionally hypermethylated (Fig. 2A). In addition, the putative promoter regions of the *Insc* long-form transcripts and the pseudogene loci (*Gm18599*) were located near or within hypermethylated regions, suggesting epigenetic silencing and transcriptional inactivity, whereas *Gm15500* was located outside of these hypermethylated regions (Fig. 2A).

**Fig. 2.**
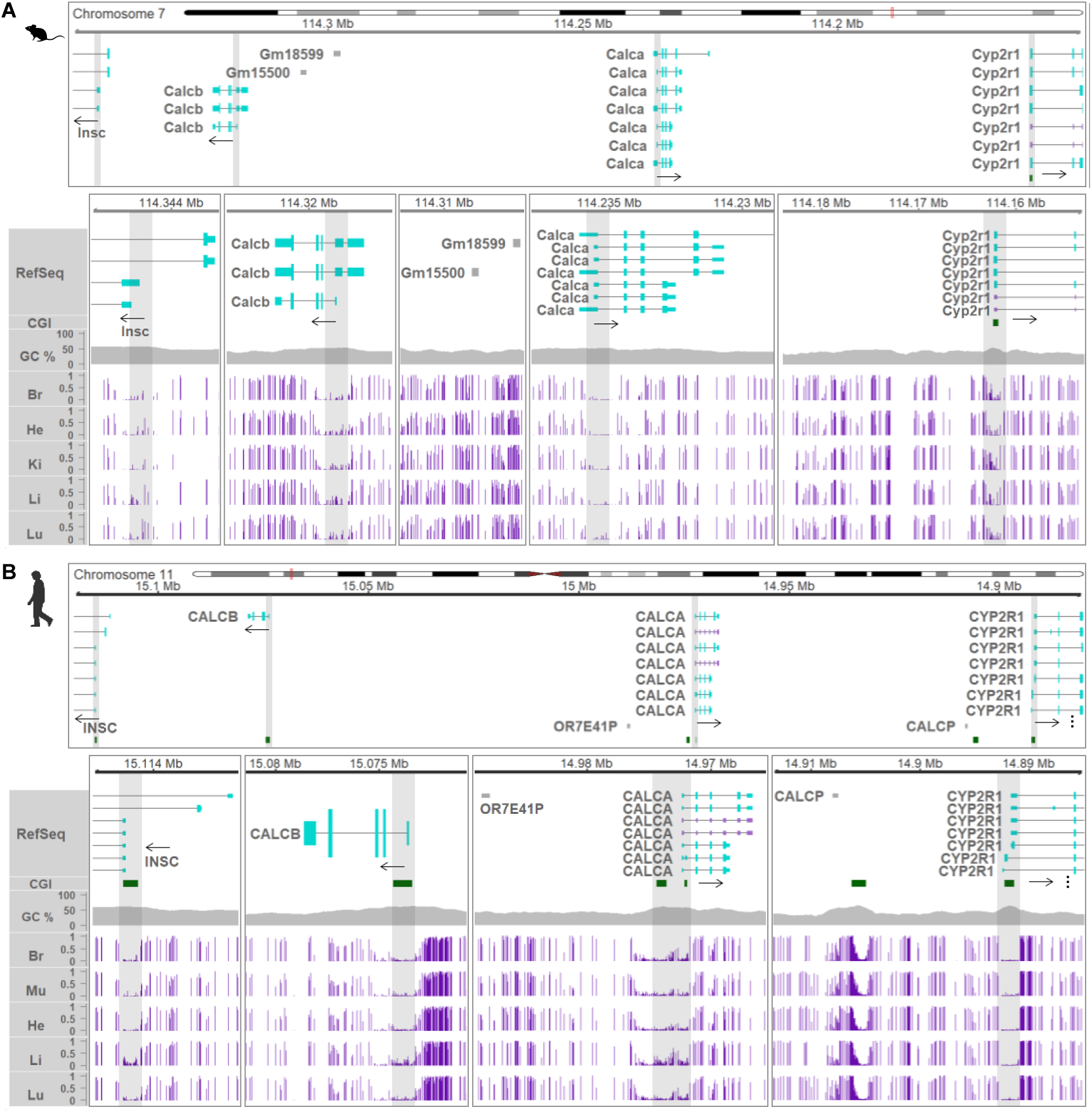
DNA methylation profiles of the mouse and human along the *Insc*/*INSC*‒*Cyp2r1*/*CYP2R1* region**. (A)** The mouse genomic interval chr7:114,350,000‒114,150,000 (reverse strand) is displayed. A zoomed view focuses on CpG methylation at putative promoter regions (gray shading) of *Insc*, *Calcb*, *Calca*, and *Cyp2r1*, as well as at the pseudogene loci *Gm15500* and *Gm18599*. **(B)** The human genomic interval chr11:15,120,000‒14,880,000 (reverse strand) is presented. A zoomed view focuses on CpG methylation at putative promoters (gray shading) of *INSC*, *CALCB*, *CALCA*, and *CYP2R1*, as well as at the pseudogene loci *OR7E41P* and *CALCP*. Transcriptional directions are indicated by arrows. Protein-coding and noncoding transcripts are shown in cyan and purple, respectively. Tall and short boxes represent translated and untranslated regions, respectively. RefSeq, NCBI RefSeq gene model; CGI, CpG island; GC%, GC content; Br, brain; He, heart; Ki, kidney; Li, liver; Lu, lung; Mu, muscle.

The order of protein-coding genes is conserved between mice and humans and, in humans, this locus is located on human chromosome 11 (cytoband 11p15.2; chr11:15.12‒14.88 Mb, reverse strand): *INSC*, *CALCB*, *CALCA*, and *CYP2R1*, with no CRSP-encoding genes present (Fig. 2B, Fig. S2). Analysis of human adult WGBS data from the NIH Roadmap Epigenomics Project revealed that all putative promoter regions, encompassing the TSS and including CpG islands with high GC content, were characteristically unmethylated or markedly hypomethylated across all analyzed tissues (brain [ectoderm], heart and muscle [mesoderm], and liver and lung [endoderm]), representing all three germ layers (Fig. 2B). Additionally, the putative promoter regions of the *INSC* long-form transcripts and pseudogenes (*OR7E41P* and *CALCP*) were situated outside unmethylated or markedly hypomethylated regions, suggesting that these loci are likely epigenetically repressed and transcriptionally inactive (Fig. 2B).

In contrast, the arrangement of protein-coding genes between *INSC* and *CYP2R1* in pigs, located on chromosome 2 (chr2:44.02‒44.37 Mb, forward strand), is not conserved relative to mice and humans (Fig. 2A,B), as *INSC*, *CALCB*, several CRSP-encoding genes (including *CALCB_1*), and *CYP2R1* are annotated, whereas the *CALCA* gene is currently absent from the annotation (Fig. 3A). In newborn pig tissues (PRJEB42772), unlike in mice and humans (Fig. 2A,B), the putative promoter regions of *CRSP3*, *CRSP-2*, *CALCB_1*, and *CRSP1-like* exhibited a partial methylation pattern (Fig. 3A). Closer examination of these promoter regions revealed partial methylation patterns distinct from the typical unmethylated or markedly hypomethylated promoter regions, with *CRSP3* showing a broader domain of partial methylation extending into the adjacent region near the first exon of the middle-form transcript (Fig. S3). On the other hand, the putative promoters of pig *INSC*, *CALCB*, and *CYP2R1* showed a typical unmethylated pattern across their promoter regions (Fig. 3A), which supports biallelic transcriptional activation, whereas the *CRSP-2* long-form promoter was located within a hypermethylated region (Fig. 3A), suggesting epigenetic silencing and transcriptional inactivation.

**Fig. 3.**
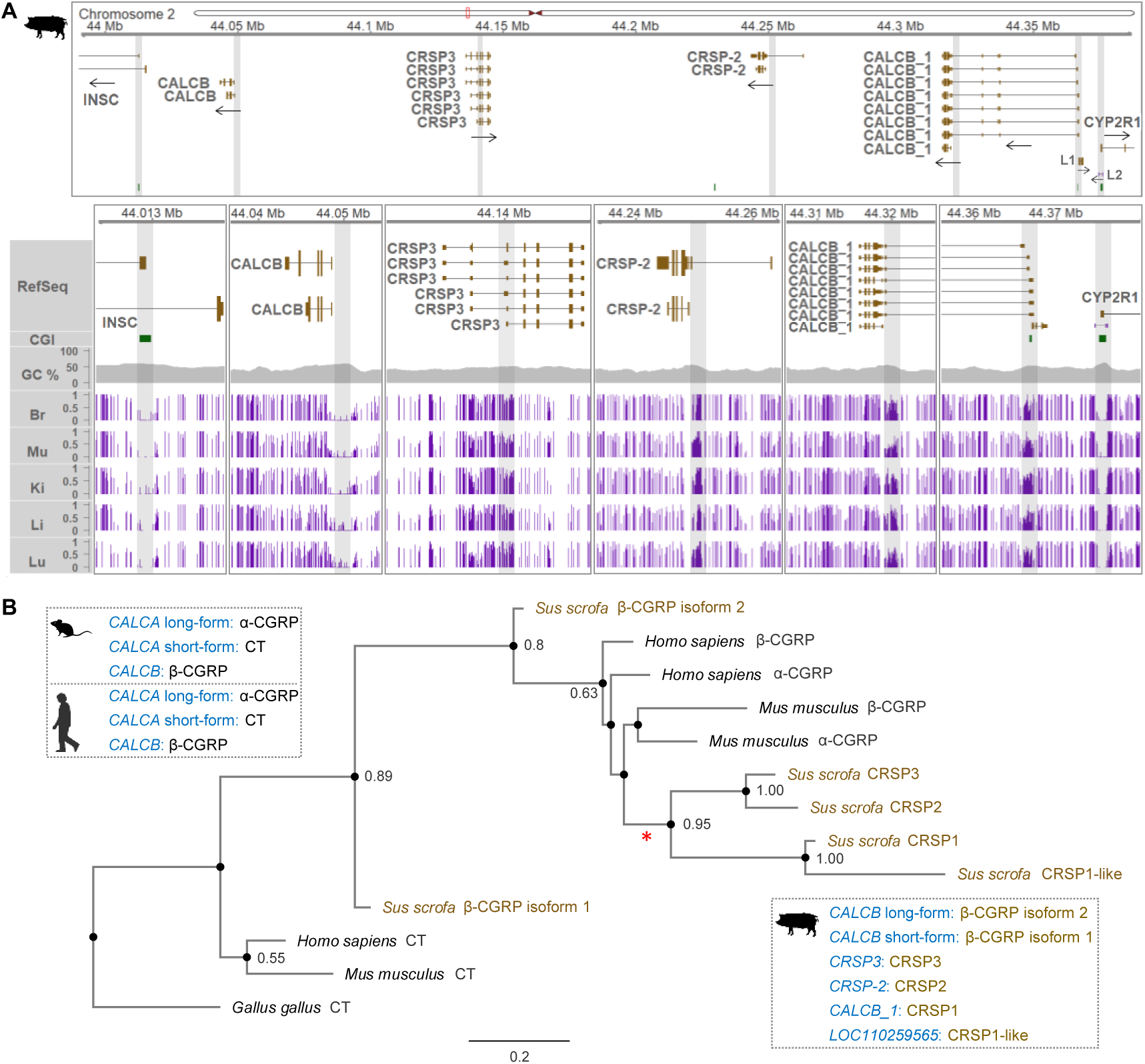
DNA methylation profiles across the porcine *INSC*‒*CYP2R1* region and phylogeny of the CT/CGRP/CRSP peptide family. **(A)** DNA methylation in newborn pig tissues (PRJEB42772) is shown across promoters and promoter-associated CpG islands withing the *CRSP* complex locus. L1, *LOC110259565* (*CRSP1-like*); L2, *LOC110259296* (uncharacterized ncRNA gene). Br, brain; Mu, muscle; Ki, kidney; Li, liver; Lu, lung. **(B)** Phylogenetic analysis of CT/CGRP/CRSP peptide family sequences from mice, humans, and pigs. Amino acid sequences of CT, α-CGRP, β-CGRP, and CRSP precursor peptides retrieved from the NCBI Gene database were used (human CT: NP_001732.1; mouse CT: NP_031613.2; human α-CGRP: NP_001365879.1; human β-CGRP: NP_000719.1; mouse α-CGRP: NP_001029126.1; mouse β-CGRP: NP_473425.2; pig accession numbers are provided in the Results section). CT from *Gallus gallus* (chicken; NP_001258893.1) was used as an outgroup. Numbers at branches represent bootstrap support values (≥ 0.50), and the scale bar indicates substitutions per site. A pig lineage-specific CRSP clade is highlighted with a red asterisk. The dashed box summarizes transcripts and their encoded amino acids.

In addition, comparative genomic views revealed that the macrosyntenic organization and local gene arrangement are largely conserved among mice, humans, and pigs, with the pig genome inverted relative to the others, across an approximately 1.2‒2.8 Mb window encompassing the *INSC*‒*CYP2R1* region (Fig. S4A‒C). Notably, the *CRSP* complex locus in pigs exhibited structural variability compared to mice and humans (Fig. S4B,C). This suggests that a microsyntenic disruption might have occurred at the porcine *CRSP* complex locus and that this locus is evolutionarily divergent among the three analyzed mammalian species (mouse, human, and pig).

In summary, comparative analyses across mice, humans, and pigs unveiled a distinct promoter methylation pattern at the *CRSP* complex locus specifically in pigs, highlighting that this locus is associated with microsyntenic rearrangement.

### Phylogeny of the CT/CGRP/CRSP peptide family supports the emergence of porcine-specific genomic imprinting of CRSP-encoding genes

With the aim of gaining further evolutionary insight into genomic imprinting, we performed phylogenetic analysis of the amino acid sequences of CT/CGRP/CRSP precursor polypeptides. Phylogenetic clustering with strong bootstrap support values (0.95–1.00) indicates that porcine CRSP2 (XP_020935894.1) and CRSP3 (XP_005661168.1), as well as CRSP1 (XP_020935854.1; encoded by *CALCB_1*) and CRSP1-like (XP_020940714.1), likely arose from gene duplication events unique to the pig lineage after its divergence from the common ancestor shared with humans and mice (Fig. 3B). In addition, two isoforms of β-CGRP (encoded by *CALCB*) in pigs show distinct evolutionary trajectories: β-CGRP isoform 1 (XP_005661167.1) clusters near the CT clade, whereas β-CGRP isoform 2 (NP_001095943.1) is nested within the CGRP/CRSP clade (Fig. 3B). This suggests isoform-specific divergence of *CALCB*-derived peptides in pigs. The phylogenetic position of CRSPs, together with the methylation status of the orthologous regions between *INSC* and *CYP2R1* and the porcine-specific imprinting of the CRSP-encoding genes, suggests that genomic imprinting at the *CRSP* complex locus may have evolved relatively recently following gene duplication in pigs.

### CRSP-encoding genes within the *CRSP* complex locus display porcine-specific imprinted monoallelic expression

Imprinted gene expression is identified by detecting consistent parent-of-origin-specific allelic expression (maternal or paternal) in crossbred offspring from initial and reciprocal crosses, irrespective of the breed pairing used in the crosses. To determine whether genes in the *CRSP* complex locus exhibit imprinted expression, we: (i) analyzed whole-genome sequencing (WGS) data from two initial and two reciprocal pig crosses (sire, dam, and F_1_ offspring) for haplotype phasing, and (ii) subsequently examined RNA-seq data from F_1_ offspring to tag reads by haplotype (haplotag) and assess parental allele-specific expression in somatic tissues (Fig. S5A).

First, for haplotype phasing, we performed combined pedigree- and read-based phasing of WGS-derived single nucleotide variants (SNVs) on chromosome 2 to generate a phased variant call format (VCF) file for each trio, using WGS data with mean sequencing depths of approximately 36× to 50× across four trios (Table S3). This approach yielded phased haplotype blocks (phase sets) for heterozygous SNVs in the *INSC*‒*CYP2R1* region (Fig. S5B; Table S3), within which the paternal and maternal haplotypes in each F_1_ offspring were resolved. As a result, 2,926 to 3,220 SNVs were phased within the longest phase sets (haplotype blocks), which spanned 779,220 to 779,378 bp across trio 1‒4 in the *INSC*‒*CYP2R1* region (Fig. S5B; Table S3).

Secondly, we evaluated RNA-seq expression of genes in the *INSC*‒*CYP2R1* region in somatic tissues of the F_1_ offspring, including brain [ectoderm], heart, kidney, skeletal muscle, and spleen [mesoderm], as well as liver and lung [endoderm], representing all three germ layers (Fig. 4; Figs. S6–S9). Across both the initial and reciprocal crosses, *CALCB* and *LOC110259565* (*CRSP1-like*) showed no detectable mRNA expression in any of the analyzed tissues of the F_1_ offspring, whereas the middle-form transcripts of *CRSP3*, which encompass the short-form region, and the short-form transcript of *CRSP-2* were highly expressed in the brain relative to other tissues (Fig. 4A,B; Fig. S6A,B and S8A,B). In contrast, in both initial and reciprocal crosses, *CALCB_1* was ubiquitously expressed across the analyzed tissues, with lower expression in skeletal muscle (Fig. 4A,B; Fig. S6A,B and S8A,B).

**Fig. 4.**
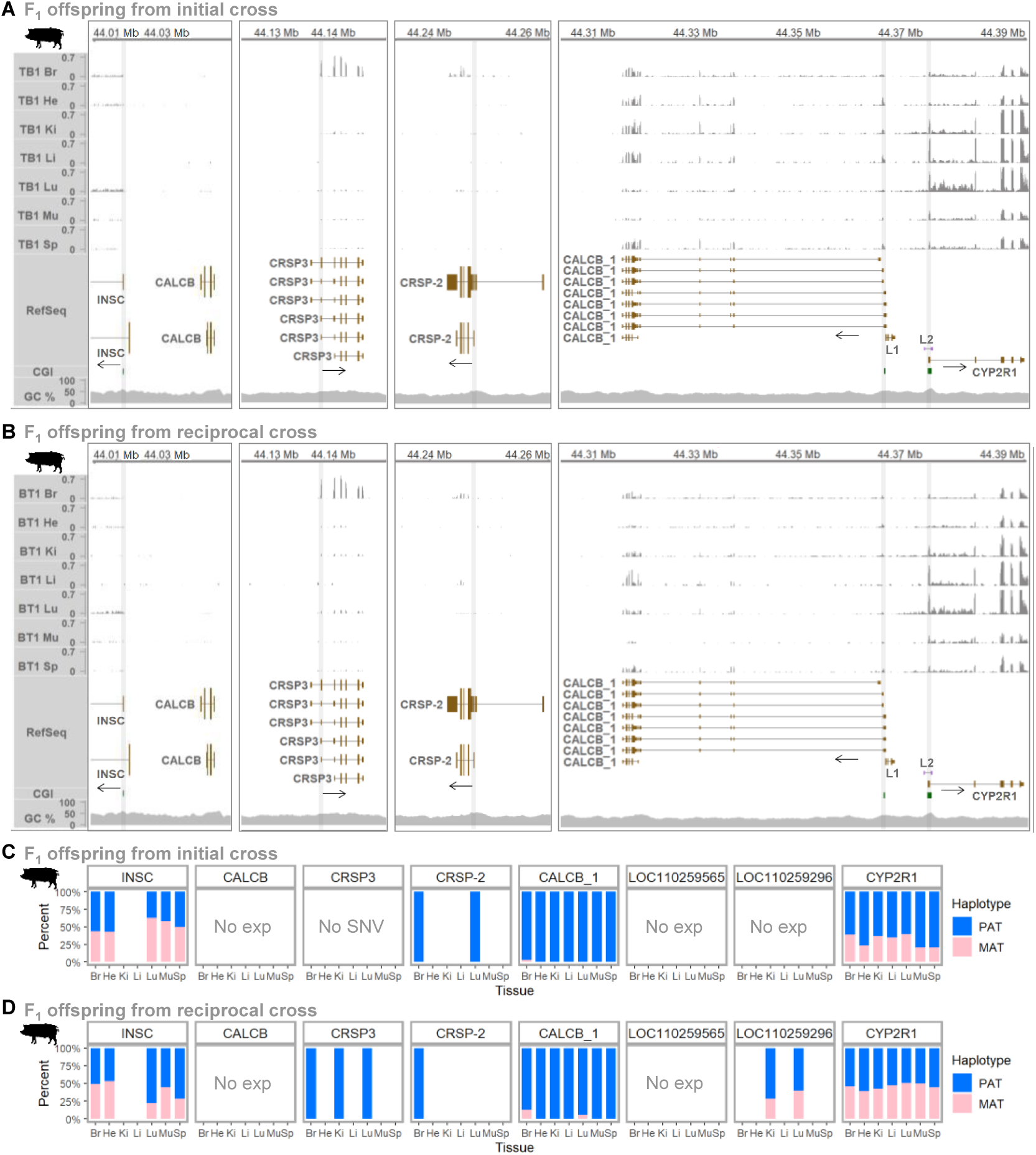
RNA expression in somatic tissues and analysis of parent-of-origin-specific (imprinted) expression. **(A, B)** RNA expression based on RNA-seq in F1 offspring from initial and reciprocal crosses of Tibetan (T) and Berkshire (B) pigs (PRJNA998604), shown as representative presentation of Fig. S6A,B (initial) and Fig. S8A,B (reciprocal). TB, F1 offspring of Tibetan sire × Berkshire dam cross (initial); BT, F1 offspring of Berkshire sire × Tibetan dam cross (reciprocal); Br, brain; He, heart; Ki, kidney; Li, liver; Lu, lung; Mu, skeletal muscle; Sp, spleen. **(C, D)** Percentages of haplotype-tagged reads for paternal (PAT) and maternal (MAT) haplotypes, including only gene–tissue combinations with >5 haplotype-tagged reads (maximum depth: 427), presented using summed counts pooled from Fig. S6C,D (initial) and Fig. S8C,D (reciprocal). L1, *LOC110259565*; L2, *LOC110259296*; No exp, no detectable RNA expression across tissues; No SNV, absence of heterozygous exonic SNVs.

Using these RNA-seq data, reads were subsequently assigned to the paternal or maternal haplotype (haplotagging; haplotype tagging) based on the phased VCF generated in the first step, enabling quantification of haplotype-specific reads. As a result, parent-of-origin-specific (paternal) expression pattern was observed for *CALCB_1* across all analyzed tissues and for *CRSP-2* in the brain and lung (Fig. 4C,D; Table S3). *CRSP3* also exhibited paternal expression in the reciprocal cross but was not detected in the initial cross due to the absence of heterozygous SNVs within exons (Fig. 4C,D; Table S3). Analysis of brain subregions from an independent individual revealed monoallelic expression pattern of *CRSP3* across all examined regions, including the cerebellum, mortor cortex, hippocampus, hypothalamus, substantia nigra, and thalamus (Fig. S10; Table S3), providing additional support for monoallelic expression of *CRSP3*. Conversely, the flanking genes *INSC* and *CYP2R1* exhibited both paternal and maternal haplotypes in both the initial and reciprocal crosses, indicating consistent biallelic expression, while *LOC110259296* (*ncRNA*) also showed both haplotypes in the reciprocal cross but was not detected the initial cross due to lack of expression in the analyzed offspring tissues (Fig. 4C,D; Table S3). Additionally, haplotype tagging was consistent with paternal monoallelic expression, as exemplified by *CALCB_1* and *CRSP-2* in the initial cross (Fig. S7) and by *CRSP3* and *CALCB_1* in the reciprocal cross (Fig. S9).

Consequently, using a haplotype-based approach, we identified parent-of-origin-specific (paternal) expression of the CRSP-encoding genes *CRSP-2* and *CALCB_1* (*CRSP1*), regardless of breed combination, indicating their imprinted paternal expression. Paternal expression of *CRSP3* was further partly supported by monoallelic expression patterns observed across pig brain subregions.

### Comparative analyses in mice reveal haplotype-resolved biallelic expression of *Calcb* and *Calca* at the orthologous locus

In comparison with porcine imprinted expression, analyses in mice were conducted to assess whether gene expression at the orthologous locus resembles or differs from that in pigs. Taking advantage of the inbred nature of mouse strains and using the C57BL/6J reference genome (GRCm39/mm39), we analyzed initial and reciprocal crosses between C57BL/6J and CAST/EiJ mice.

As an initial step, instead of performing WGS-based SNV phasing, we leveraged strain-specific variants from the Mouse Genomes Project (MGP; REL2021) to generate phased variant sets for allele-specific expression analysis. Reference alleles were assigned to C57BL/6J and alternative alleles to CAST/EiJ, enabling inference of parental haplotypes in F_1_ offspring from initial and reciprocal crosses. This strategy yielded computationally phased VCFs for the offspring from each cross that distinguish paternal and maternal alleles for downstream haplotype-resolved expression analysis.

Next, gene expression was examined using RNA-seq data from the initial and reciprocal crosses (accession SRP020526 [23]) prior to haplotype-tagging. In the initial cross, *Calcb* showed high expression in the duodenum, whereas *Calca* was highly expressed in the hypothalamus (Fig. S11A). In the reciprocal cross, a similar expression pattern was observed, although *Calca* expression in the hypothalamus was reduced (Fig. S11B). In both the initial and reciprocal crosses, *Insc* showed high expression in the brain (cerebellum and whole brain), whereas *Cyp2r1* exhibited consistently low expression across the analyzed tissues (Fig. S11A,B). Additionally, the pseudogene *Gm15500* was ubiquitously expressed, and *Gm18599* was not detectable (Fig. S11A,B).

Subsequently, haplotagging on the RNA-seq reads detected both paternal and maternal haplotypes consistently in both the initial and reciprocal crosses (Fig. 5A,B; Fig. S11C,D; Table S4), indicating a biallelic expression pattern for *Insc*, *Calcb*, and *Calca* across the analyzed tissues (cerebellum, duodenum, embryonic brain, hypothalamus, lung, and whole brain), with a few tissues omitted due to low coverage. In addition, visualization of haplotagged RNA-seq reads in IGV further supported biallelic expression in both the initial and reciprocal crosses, as exemplified by *Calcb* and *Calca* (Figs. S12 and S13). To evaluate *Cyp2r1* expression, we analyzed an additional dataset that showed high expression of *Cyp2r1* in the liver (Fig. S14A,B). Haplotagging detected both paternal and maternal haplotypes, indicating biallelic expression of *Cyp2r1* (Fig. 5A,B; Fig. S14C,D; Table S4), consistent with the patterns observed for the other genes (*Calcb*, *Calca*, and *Insc*) .

**Fig. 5.**
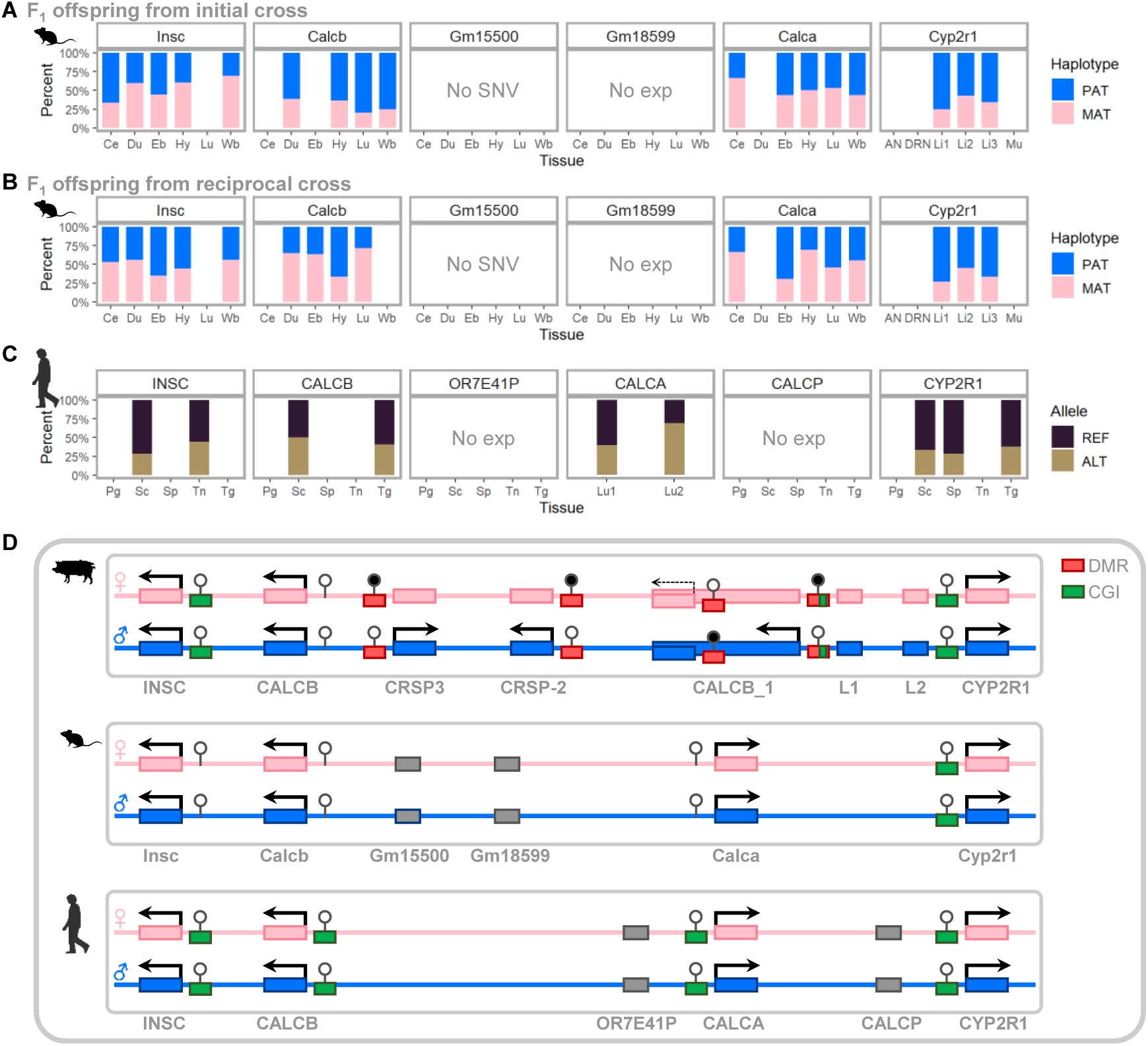
Allelic expression at the *INSC*–*CYP2R1* locus across species. **(A, B)** Percentages of haplotype-tagged reads representing paternal (PAT) and maternal (MAT) haplotypes in mice. Ce, cerebellum; Du, duodenum; Eb, embryonic brain; Hy, hypothalamus; Lu, lung; Wb, whole brain; AN, arcuate nucleus; DRN, dorsal raphe nucleus; Li, liver; Mu, muscle. **(C)** Percentages of reference (REF) and alternative (ALT) allele counts in humans. Pg, prostate gland; Sc, sigmoid colon; Sp, spleen; Tn, tibial nerve; Tg, thyroid gland; Lu, lung. No SNV, absence of CAST/EiJ strain-specific SNVs; No exp, no detectable RNA expression across tissues. **(D)** Schematic representation of biallelic and parent-of-origin-specific expression at the *INSC*–*CYP2R1* locus in pigs, mice, and humans. Maternal and paternal alleles are shown in pink and blue, respectively. Arrows denote transcriptional direction. The short-form *CALCB_1* transcript in pigs is depicted with a dashed arrow to reflect low expression. Gray boxes represent pseudogenes. Filled and unfilled circles denote methylated and unmethylated regions, respectively.

Overall, haplotype-resolved allele-specific expression analysis revealed that *Calcb* and *Calca* are biallelically expressed across somatic tissues in mice in both the initial and reciprocal crosses, in contrast to the cluster of paternally expressed genes at the orthologous locus in pigs.

### Analyses in humans reveal biallelic expression of *CALCB* and *CALCA* at the orthologous locus

Unlike in pigs and mice, where experimental initial and reciprocal crosses are used, monoallelic and biallelic expression patterns in humans were assessed using matched genomic and RNA expression data from the same individuals. Using human ENCODE raw WGS data, informative heterozygous SNVs were identified in the last exon of the *CALCB* gene in two adult individuals (donor IDs: ENCDO451RUA and ENCDO845WKR) (Fig. S15A).

Gene expression was then examined across multiple tissues (prostate gland, sigmoid colon, spleen, tibial nerve, and thyroid gland) from these individuals. In both individuals, the short isoform of *CALCA* was expressed specifically in the thyroid gland; *CALCB* in the sigmoid colon and thyroid gland; *INSC* in the sigmoid colon and tibial nerve; and *CYP2R1* in all analyzed tissues, with higher expression levels in the first individual (ENCDO451RUA) than in the second (ENCDO845WKR) (Fig. S15B,C).

In the first individual, a biallelic expression pattern of *CALCB* was consistently observed in two tissues (sigmoid colon and thyroid gland) (Fig. 5C; Fig. S15D,F; Table S5), whereas its expression was too low in the second individual to determine this pattern (Fig. 5C; Fig. S15E; Table S5). Additionally, *INSC* in the first individual and *CYP2R1* in the second individual also showed biallelic expression tendencies (Fig. 5C; Fig. S15D,E).

Furthermore, in human lung exome data, heterozygous SNVs were identified in exon 4 of the *CALCA* gene in two adult individuals (Ind 9 and Ind 30) (Fig. S16A). Gene expression in lung RNA-seq data were then examined, revealing expression of the long isoform of *CALCA* in both individuals (Fig. S16B,C). In both cases, reads carrying the reference and alternative alleles were detected in lung tissues, indicating a biallelic expression pattern (Fig. 5C; Fig. S16D,E).

In addition to the biallelic expression observed in mice, biallelic expression of human *CALCB* and *CALCA* was also detected, further supporting the conclusion that the paternally expressed gene cluster at the orthologous locus is specific to pigs. In summary, the comparative schematic view depicts the presence of maternally imprinted and paternal expressed genes in pigs, including low expression of the *CALCB_1* short-form, and the absence of such imprinting in mice and humans at the locus between *INSC* and *CYP2R1* (Fig. 5D), highlighting porcine-specific imprinting at this locus.

### Oocytes-derived alternative transcription may underlie methylation imprints at the *CRSP* complex locus

To investigate potential mechanisms underlying clustered imprinting at the *CRSP* complex locus in pigs, we integrated evolutionary, developmental, and transcriptional perspectives using a multi-omics approach that included transcriptomes, ChIP-seq enrichment, and DNA methylomes. As described above for phylogenetic analysis, this locus shows strong evidence of gene duplication specific to the pig lineage. In addition, because oogenesis is highly transcriptionally dynamic, alternative transcripts from these duplicated genes may be generated during oocyte development. In this regard, long terminal repeat (LTR) sequences, which have been shown to initiate transcription during oogenesis, were examined at the porcine *CRSP* complex locus.

In fully grown oocytes (FGOs) from pigs, the long-form *CRSP3* transcripts was expressed (Fig. 6A) and initiated adjacent to a solo LTR (MLT1D), a member of the ERVL (endogenous retrovirus group L)–MaLR (mammalian apparent LTR retrotransposon) family that can function as an oocyte-specific promoter or enhancer [24] (Fig. 6B). This solo LTR, identified through the UCSC Genome Browser, is 106 bp in length and located on the positive strand (+), consistent with the *CRSP3* transcriptional orientation (Fig. 6B). Because the H3K4me3 enrichment (a hallmark of active promoters) in oocytes lies downstream of the solo LTR and overlaps the TSS and first exon of the *CRSP3* long-form transcript (Fig. 6C), the upstream LTR may function as an enhancer rather than a promoter. At later developmental stages (4-cell, 8-cell, and blastocyst), the promoter mark shifted to regions near the TSS and first exon of the middle- and short-form transcripts (Fig. 6C), suggesting stage-specific expression of these isoforms. Additionally, WGBS data revealed full methylation in oocytes, unmethylation or hypomethylation in sperm, and partial methylation in somatic tissue (muscle) near the putative promoters and TSS of the middle- and short-form transcripts (Fig. 6D; Fig. S17), suggesting imprinted expression of these isoforms in somatic tissue.

**Fig. 6.**
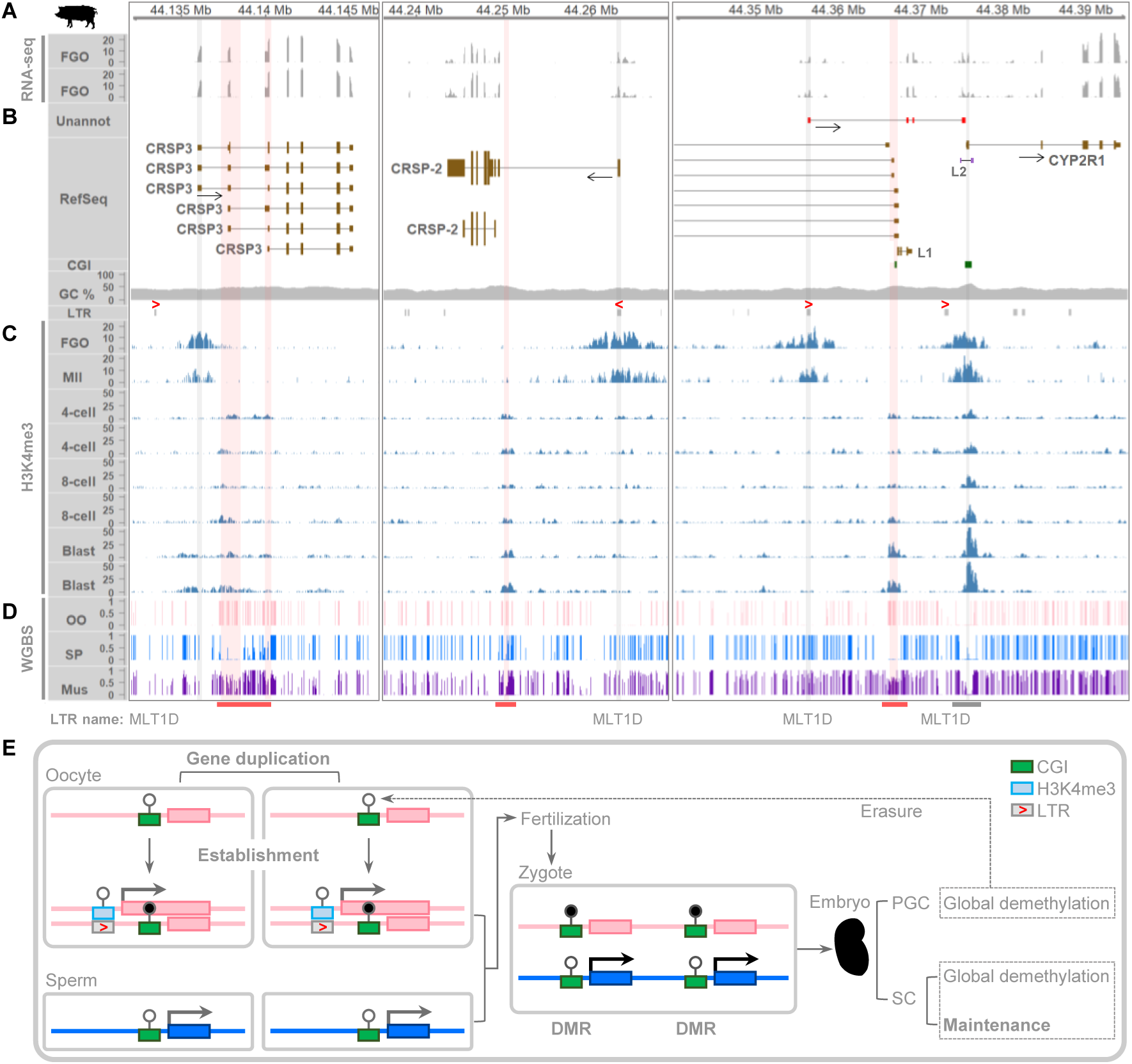
Transcription-derived methylation imprinting in intragenic regions at the *CRSP* complex locus. **(A)** RNA-seq read coverage profiles retrieved from porcine fully grown oocytes (GSE163709). The coverage values were normalized equivalently to transcripts per million (TPM). **(B)** An unannotated transcript (Unannot) and annotated transcripts from the NCBI RefSeq annotation (RefSeq). The transcriptional directions of expressed transcripts are denoted by arrows. L1, *LOC110259565* (*CRSP1-like*); L2, *LOC110259296* (uncharacterized ncRNA gene); CGI, CpG island; CG%, GC content; and LTR, long terminal repeat adapted from the UCSC Genome Browser. The orientation of selected LTRs is indicated by red arrowheads, and the name of these LTRs (MLT1D) is shown at the bottom of the plot. **(C)** Enrichment patterns of H3K4me3 histone modifications during pig development. ChIP-seq data were normalized to 1× genomic coverage (reads per genome coverage, RPGC). FGO, fully grown oocyte; MII, metaphase II oocyte (GSE163620); 4-cell, 4-cell stage embryo; 8-cell, 8-cell stage embryo; Blast, blastocyst (GSE163709). **(D)** WGBS data from oocytes (OO) and sperm (SP) (GSE143850), and from somatic muscle tissue (Mus) of a 120-day-old Landrace pig (GSE157045). Regions marked with red and grey bars are enlarged in Fig. S19. **(E)** Schematic representation of recurrent imprinting of CRSP-encoding genes. Establishment, DNA methylation establishment; PGC, primordial germ cell; SC, somatic cell; Erasure, erasure of DNA methylation imprints; Maintenance, maintenance of DNA methylation imprints.

For the *CRSP-2* gene, the long-form transcript was expressed in FGOs (Fig. 6A), and an MTL1D solo LTR (476 bp) located on the negative strand (–) overlaps the TSS and first exon of this long-form transcript (Fig. 6B). Because this LTR also coincides with strong H3K4me3 enrichment in oocytes (Fig. 6C), it likely functions as an active promoter for the long-form during the oocyte stage. Consistently, the putative promoter and TSS of the long-form are unmethylated in oocytes, suggesting active transcription, whereas they are fully methylated in sperm, indicating transcriptional repression (Fig. 6D). At later developmental stages (4-cell, 8-cell, and blastocyst), H3K4me3 enrichment shifts toward the TSS of the *CRSP-2* short-form transcript (Fig. 6C), indicating that the long-form promoter becomes inactive outside the oocyte stage, whereas the short-form promoter becomes active. Additionally, WGBS data showed full methylation in oocytes, unmethylation in sperm, and partial methylation in somatic tissue (muscle) near the short-form TSS (Fig. 6D; Fig. S17). Thus, this promoter transition, together with the somatic half-methylation pattern, may underlie the emergence of imprinted expression of the *CRSP-2* short-form.

Regarding *CALCB_1*, an unannotated transcript was detected in FGOs (Fig. 6A; Fig. S18). This transcript spans the first exons of the *CALCB_1* long-form isoforms but is oriented in the opposite (forward) direction, as indicated by the orientation of the splice donor and acceptor sites (Fig. S19), and terminates immediately upstream of *CYP2R1* (Fig. 6A; Fig. S18). Another MLT1D solo LTR (347 bp; + strand) overlaps the TSS and first exon of this unannotated transcript (Fig. 6B). This LTR also coincides with H3K4me3 enrichment in oocytes (Fig. 6C), and the putative promoter and first exon are unmethylated in oocytes but fully methylated in sperm, indicating that this LTR functions as an oocyte-specific promoter that initiates transcription in oocytes but not in sperm (Fig. 6D). In later zygotic developmental stages, H3K4me3 enrichment shifts from the promoter of the unannotated transcript toward the first exon region of *CALCB_1*, with stronger enrichment observed in blastocysts. At the same time, the oocyte-specific H3K4me3 signal for the unannotated transcript TSS disappears, suggesting that this transcript is no longer expressed at these later stages (Fig. 6C). In addition, WGBS revealed full methylation in oocytes, unmethylation in sperm, and partial methylation in muscle at the first exon regions of *CALCB_1* and the associated CGI (Fig. 6D; Fig. S17), consistent with an imprinted pattern. Together, the promoter and transcript transition, along with somatic partial methylation, support the establishment and maintenance of imprinted expression of *CALCB_1*. In summary, the unannotated overlapping transcript spanning the *CALCB_1* promoter CGI may contribute to oocyte-specific CGI methylation, followed by promoter transition and partial somatic methylation, thereby supporting the establishment and maintenance of imprinted expression of the *CALCB_1* long-form.

*CYP2R1* was also expressed in FGOs (Fig. 6A,B), but no shift in the genomic position of the H3K4me3 enrichment was observed in the *CYP2R1* promoter CGI region (Fig. 6C), which remained unmethylated in oocytes, sperm, and somatic tissue (Fig. 6D; Fig. S17), consistent with a lack of imprinting at *CYP2R1*. Therefore, unlike *CYP2R1*, the CRSP-encoding genes exhibit recurrent imprinting that may have arisen following gene duplication events (Fig. 6E). This recurrent pattern is proposed to be coupled with transcription-derived imprinting at the *CRSP* complex locus (Fig. 6E). In conclusion, overlapping transcription in oocytes, together with zygotic promoter shifting, may contribute to the repeated intragenic imprinting observed at the *CRSP* complex locus.

## Discussion

In this study, we report the identification of a novel imprinted *CRSP* complex locus that appears to be unique to pigs and is absent in mice and humans. Although approximately 200 imprinted genes have been reported in mice and humans, with about 60 shared between the two species [9], several orthologous loci are not imprinted in domesticated animals [11]. This highlights species-or lineage-specific imprinting patterns (e.g., rodent vs. primate vs. artiodactyl) and suggests, as supported by recent studies [25, 26], that additional livestock-specific imprinted genes may exist beyond those currently identified but remain incompletely characterized. In this context, our findings expand the current understanding of porcine-specific imprinting. Notably, the *CRSP* complex locus spans approximately 240 kb, which is about four times narrower than some of the largest known imprinted domains that can extend up to ∼1 Mb [27, 28]. Additionally, no evident chromosomal rearrangement was detected within the 2.8-Mb flanking region in pigs compared with mice and humans, except for the *CRSP* complex locus itself. These observations suggest that the microsyntenic disruption in pigs may have accompanied the locus-specific emergence of genomic imprinting, potentially through gene duplication events that gave rise to the multiple CRSP-encoding genes. The evolutionary persistence of these duplicated genes might reflect the dosage-sharing hypothesis [29], which proposes that duplicated genes are down-regulated so that their combined expression level matches that of the ancestral single-copy gene. This down-regulation might be achieved by genomic imprinting at the *CRSP* complex locus, where silencing of one parental allele serves to offset the increased dosage associated with gene duplication.

Germline DMRs are established in the gametes and maintained after fertilization by escaping post-fertilization epigenetic reprogramming, and they persist in somatic tissues throughout the lifetime (primary DMRs), although a rare case of postnatal loss of imprinting has been reported at the *Dlk1* locus in neurogenic niche cells in mice [30, 31]. Once germline DMRs are established, methylation maintenance involves ZFP57, which recognizes the TGCCGC hexanucleotide motif and recruits the KAP1/TRIM28 complex together with the DNA methyltransferase DNMT1 to protect both maternal and paternal methylation imprints during the wave of genome-wide DNA demethylation in the early embryo [22, 32]. This consensus motif is also recognized by ZNF445/ZFP445, another key regulator factor of imprinting maintenance [33, 34], which is expressed from the porcine oocyte stage [35]. In the current study, the porcine *CRSP* complex locus displayed the canonical pattern of germline DMR persistence across embryonic, neonatal, and adult tissues, and the enrichment of TGCCGC motifs within these regions supports their capacity for methylation maintenance. In contrast, the putative promoter regions, including promoter-associated CpG islands, are typically unmethylated under normal physiological conditions [36, 37], as demonstrated by methylation patterns at non-imprinted regions within the *INSC*–*CYP2R1* locus in pigs, mice, and humans. In addition, for the porcine embryonic stage, our use of parthenogenetic embryos is based on the allelic differences between controls and parthenogenetic embryos that provide a clear parent-of-origin contrast: maternally methylated DMRs retain full or hypermethylated states, whereas paternally methylated DMRs show unmethylated or hypomethylated patterns in parthenogenetic embryos, with control embryos exhibiting the expected half methylation pattern for both classes of DMRs. This approach could be implemented without the use of androgenetic (bipaternal) embryos, which are considerably more difficult to generate in pigs [38].

Using pedigree information and genomic sequences from the sire, dam, and offspring, the two alleles at heterozygous SNV sites in the offspring can be phased into paternal and maternal haplotypes [39]. Combining pedigree-based phasing, which provides chromosome-scale haplotypes, with read-based phasing, which refines local haplotypes using sequencing reads spanning multiple variants, can improve phasing accuracy and allele-specific expression detection, particularly in regions where parental genotypes are locally uninformative (i.e., both parents are heterozygous) [40]. This integrated strategy provides more complete and contiguous phase sets than either method alone [40], thereby enhancing detection of imprinted expression of CRSP-encoding genes in pigs. Additionally, the expression patterns observed in humans are consistent with true biallelic expression rather than RNA editing, because the same heterozygous SNVs detected in WGS/exome data are also present in the RNA-seq reads. Moreover, canonical A-to-I RNA editing results in A-to-G changes in RNA (inosine being read as guanosine during sequencing) [41], which does not apply to the current study, as the variants observed are G/T and C/T rather than A/G mismatches. Likewise, in the mouse F_1_ offspring, the A/G pattern reflects true heterozygosity and biallelic transcription rather than RNA editing, as the A and G alleles observed in RNA-seq correspond to the homozygous A/A and G/G genotypes of the C57BL/6J and CAST/EiJ parental strains, respectively. Moreover, in pigs, although one F_1_ offspring had an A/G genotype at the DNA level, only the A allele was detected in RNA-seq. Similarly, in brain subregions from a pig individual heterozygous for A/G, RNA-seq showed predominant expression of the A allele for *CRSP3*. Together, these observations exclude the possibility of A-to-I (G) RNA editing and support imprinted monoallelic expression. With respect to gene expression, *CALCB_1* and *LOC110259565* (*CRSP1-like*) share a bidirectional promoter region that could, in principle, regulate imprinted expression in both directions; however, expression of *LOC110259565* was not detected, whereas *CALCB_1* was expressed, suggesting that transcriptional regulatory elements (e.g., enhancers and anti-repressing elements) are active only for *CALCB_1*. In addition, expression of the *CALCB_1* short-form transcript, which contains a non-overlapping region within the first exon, was observed to be low.

LTR retrotransposons, which characteristically contain a 5′ LTR, internal retroviral coding regions, and a 3′ LTR, constitute 7.56% of the pig genome, with endogenous retroviruses (ERVs) representing the most prominent subtype and accounting for 7.43% [42, 43]. In addition to these ERVs, solo-LTRs (recombination-derived single LTR remnants lacking the internal retroviral coding region) are also abundant, with an average of ∼5,630 copies reported per pig genome [43]. MLT1D is a family of LTR elements derived from an ancient ERVL–MaLR retrotransposon, and based on their sizes (106–478 bp) and the absence of any ‘-int’ annotations, which mark internal retroviral sequences, the MLT1D elements identified in this study are regarded as solo-LTRs. In contrast, full-length ERVs in pigs are typically 8.5–11 kb in length [42]. Eukaryotic transposable elements (TEs), including LTR retrotransposons, can integrate at diverse genomic locations [44], and repeated MLT1D solo-LTRs within or near the putative promoter regions at the porcine *CRSP* complex locus may be associated with ancestral locus-specific gene duplication events that contributed to the evolution of the imprinting cluster.

Species-specific distributions of LTRs at orthologous loci contribute to divergent transcription in mammalian oocytes, and LTR-initiated transcription has been reported to induce intragenic CGI methylation and its maintenance, thereby establishing maternally methylated DMRs in humans, mice, and pigs [25, 26, 45–47]. Specifically, the resulting gene-body methylation at CGIs embedded within oocyte-specific transcripts precedes DNMT3A/3L-dependent *de novo* DNA methylation during oogenesis [46] and is subsequently preserved after fertilization through ZFP57/KAP1-mediated maintenance methylation at imprinted loci [22]. Our results strongly suggest that this mechanism also operates at the porcine *CRSP* complex locus.

The imprinted expression of the CRSP-encoding genes may influence calcitonin receptor signaling in the CNS and peripheral tissues, thereby potentially modulating calcium homeostasis. Whether this imprinting aligns with the parental conflict theory, which posits that paternally expressed genes promote growth whereas maternally expressed genes are related to reduced growth [3, 4], requires further investigation to fully clarify the role of these imprinted genes in pig development and growth. Future studies may include analyses of single nucleotide variants or other mutations in regulatory or coding regions associated with growth traits, as well as investigations examining the consequences of loss of imprinting leading to biallelic expression. From an applied perspective, parental breeding values for sires and dams can be estimated more accurately by incorporating imprinting effects using information on the parental origin of alleles at imprinted loci [8]. Therefore, identifying new imprinted genes and defining the scope of genomic imprinting in pigs are essential for improving the accuracy of genomic selection and enhancing swine productivity.

## Conclusions

By integrating comparative genomics, epigenomics, and transcriptomics approaches, we revealed that the porcine *CRSP* complex locus harbors parent-of-origin-specific differentially methylated regions and imprinted genes that are not conserved in humans or mice. These results highlight the *CRSP* complex locus as a novel porcine-specific imprinted domain and provide a foundation for future studies of lineage-specific genomic imprinting. Further analyses provided mechanistic insight into how gene duplication may have contributed to the emergence of this imprinted domain, warranting functional characterization of the *CRSP* complex locus.

## Methods

### Sample preparation

The procedures for *in vitro* maturation of pig oocytes and for producing parthenogenetically activated (PA) embryos followed our previous reports [48, 49]. Briefly, ovaries from LYD (Landrace x Yorkshire x Duroc) pigs were collected from a local slaughterhouse, and oocytes were matured and activated *in vitro*. Following activation, PA embryos were transferred into the oviducts of two 12-month-old LYD surrogate gilts. Fertilized control (CN) embryos were obtained by naturally mating two LYD gilts with boars twice at 6-h interval at the onset of estrus. PA and CN embryos were recovered from euthanized surrogates and gilts, respectively, on day 21 after the onset of estrus, prior to the appearance of morphological changes in parthenogenetic embryos. Based on our previous histological analyses demonstrating normal PA embryo development up to day 26 [50], morphologically intact embryos were isolated from placental tissue and cryopreserved in liquid nitrogen until further analysis.

### Whole genome bisulfite sequencing

WGBS methylomes were generated independently for each individual in the CN and PA groups, as previously reported [25]. Genomic DNA was isolated from whole embryos (n = 3 per group) and used for WGBS library preparation with the Accel-NGS Methyl-Seq DNA Library Kit (Swift Biosciences). Bisulfite conversion efficiency exceeded 99%, as estimated using lambda phage spike-in controls. Libraries were sequenced as 151-bp paired-end reads on an Illumina HiSeqX platform. Raw reads were trimmed to remove adapter sequences and filtered to exclude low-quality reads using Trim Galore (v0.6.6) with default parameters, resulting in more than 800 million cleaned reads per sample. Reads were aligned to the pig reference genome (Sscrofa11.1) using Bismark (v0.25.1) with default parameters. After deduplication, mean alignment depth was ∼35× per sample. Methylation ratios for all CpG cytosines were extracted using the Bismark methylation extractor with default settings for paired-end reads. CpG sites covered with at least three reads in each sample were further analyzed. Median CpG coverage was 10–11× per sample. DMRs were identified using metilene (v0.2-8) with default parameters and the following criteria: a maximum distance of 300 bp between adjacent CpGs, at least 10 CpGs per region, and a false discovery rate (FDR) < 0.05. To visualize methylation levels, the R/Bioconductor package Gviz (v1.44.0) was used.

### Motif analysis

Motif scanning was performed using the FIMO (Find Individual Motif Occurrences) tool from the MEME Suite with the consensus ZFP57 binding motif (TGCCGC; MA1583.2) obtained from the JASPAR database. Motif occurrences with q-value < 0.05 were considered significant. The motif logo was generated using the R package universalmotif (v1.20.0).

### Phylogenetic tree construction

Multiple amino acid sequence alignments were performed using the MUSCLE algorithm implemented in MEGA (v11), and a maximum-likelihood tree was constructed using the JTT substitution model, which was identified as the best-fitting model based on the lowest Bayesian information criterion (BIC) score among the tested models, with 500 bootstrap replicates. The resulting Newick file, containing boostrap support values (0.00–1.00) and branch lengths, was imported into FigTree (v1.4.4) for tree visualization and editing.

### Public data processing

Raw sequencing reads from publicly available data (Table S6) were used for WGS, RNA-seq, and ChIP-seq analyses and aligned to the reference genomes Sscrofa11.1 (pig), GRCm39 (mouse), and GRCh38 (human), or genomic coordinates from earlier assemblies were converted to these references using the UCSC liftOver tool.

### Whole-genome sequencing processing and variant calling

Raw paired-end WGS data from pigs (100 bp; PRJNA998604) were retrieved and subjected to quality assessment and adapter trimming using Trimmomatic (v0.39). Cleaned reads were aligned to the pig reference genome (Sscrofa11.1) using BWA-MEM (v0.7.17), followed by duplicate marking. Variant calling was performed using pbrun haplotypecaller with GPU acceleration (NVIDIA Parabricks v4.4.0). Individual GVCFs generated for each sire, dam, and offspring were combined for each trio using GATK CombineGVCFs (GATK v4.6.0.0) and jointly genotyped with pbrun genotypegvcf (NVIDIA Parabricks v4.4.0). These Parabricks modules are functionally equivalent to the corresponding steps in the GATK Best Practices workflow.

### RNA-seq analysis

Paired-end RNA-seq data were retrieved as described above. After adapter trimming and quality filtering, cleaned reads were aligned to the respective reference genomes using the STAR aligner (v2.7.11b) with default parameters. BAM files of aligned reads were processed for duplicate marking and quality filtering using SAMtools. BAM files were normalized using bamCoverage from deepTools (v.3.5.6), and read coverage was visualized using the Gviz package.

### Phasing and haplotagging in pigs

Trio VCFs generated from WGS data were phased using WhatsHap phase (v2.3) in pedigree mode, which integrates pedigree structure and read-level information to produce phased VCFs with haplotype-resolved, parent-of-origin-assigned variant calls. The phased offspring VCFs from each trio were then used to haplotag RNA-seq reads with the WhatsHap haplotag function, assigning HP tags (HP:1 for paternal and HP:2 for maternal) to indicate haplotype origin within each phase set.

### Pseudo-phasing and haplotagging in mice

Strain-specific SNVs were obtained from the Mouse Genomes Project (MGP; REL2021), which uses the C57BL/6J reference genome (GRCm39/mm39). Informative single nucleotide polymorphisms (SNPs) were extracted with extract_snps.py [51], retaining high-quality REF and ALT sites corresponding to C57BL/6J and CAST/EiJ, respectively. For initial and reciprocal crosses, parental haplotypes in F_1_ offspring were computationally inferred as 0|1 or 1|0, respectively, generating pseudo-phased VCFs. These VCFs were subsequently used to haplotag RNA-seq reads using WhatsHap haplotag for allele-specific expression analysis.

### ChIP-seq analysis

Raw H3K4me3 ChIP-seq data were retrieved and trimmed and filtered using Trimmomatic with default settings. Cleaned reads were aligned to the pig reference genome using the BWA-MEM aligner with default parameters. Aligned reads were deduplicated and quality-filtered using SAMtools. Read coverage was normalized to 1× depth (reads per genomic content, RPGC) using bamCoverage from deepTools with the parameters --binSize 10 and --smoothLength 15. Peaks were visualized along genomic coordinates using the Gviz package.

## Declarations

## Acknowledgments

We thank the Ohio Supercomputer Center (OSC) for providing computational resources and support for this study.

## Authors’ Contributions

JA, SH, and KL conceived the project. JA developed and applied next-generation sequencing data analysis workflows. ISH and MRP contributed to parthenogenesis. JA, ISH, MRP, and ICC contributed new reagents and analytic tools. JA and KL contributed to the analysis of omics data. JA and KL drafted the manuscript. JA and KL reviewed and edited the manuscript. SH and KL contributed to funding acquisition. All authors read and approved the final manuscript.

## Availability of data and materials

All processed data are provided in the article and Supporting Information. The WGBS datasets generated in this study have been deposited in the Gene Expression Omnibus (GEO) under accession number GSE195528. Publicly available data were downloaded from the NCBI GEO repository unless otherwise stated and listed in Table S6.

## Ethics approval and consent to participate

All animal procedures were approved by the Institutional Animal Care and Use Committee of the National Institute of Animal Science, Rural Development Administration (RDA), Korea (NIAS2015-670).

## Consent for publication

Not applicable

