## Supplementary figures for "A novel imprinting cluster at the porcine *CRSP* complex locus defines a species-specific imprinted domain"

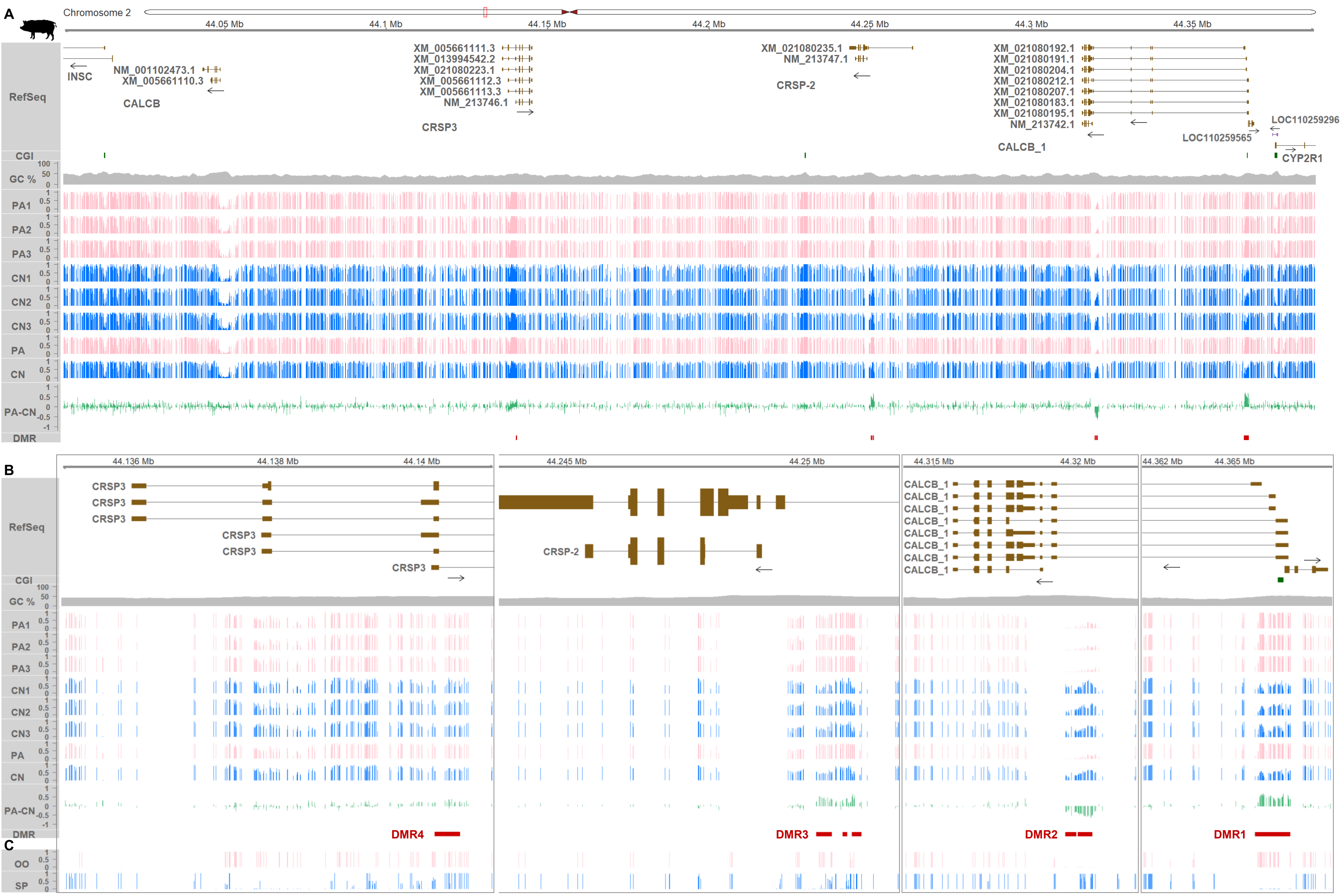


**Fig. S1. DNA methylation landscape at the porcine *CALCB–CRSP* locus. (A)** CpG methylation landscape and DMRs between parthenogenetically activated (PA) and control (CN) embryos across chr2:44,000,000–44,388,000 (*Sscrofa 11.1*; between *INSC* and *CYP2R1*), shown with NCBI RefSeq gene models. Triplicate samples for PA and CN embryos are shown, along with mean methylation levels (PA, CN) and their differences (PA – CN). **(B)** Additional gene-centered methylation profiles that complement Fig. 1B. **(C)** Germline CpG methylation in porcine oocyte (OO) and sperm (SP) across the same interval (GEO:GSE143850). Two additional unmethylated regions are shaded, beyond those highlighted in Fig. 1B–C.

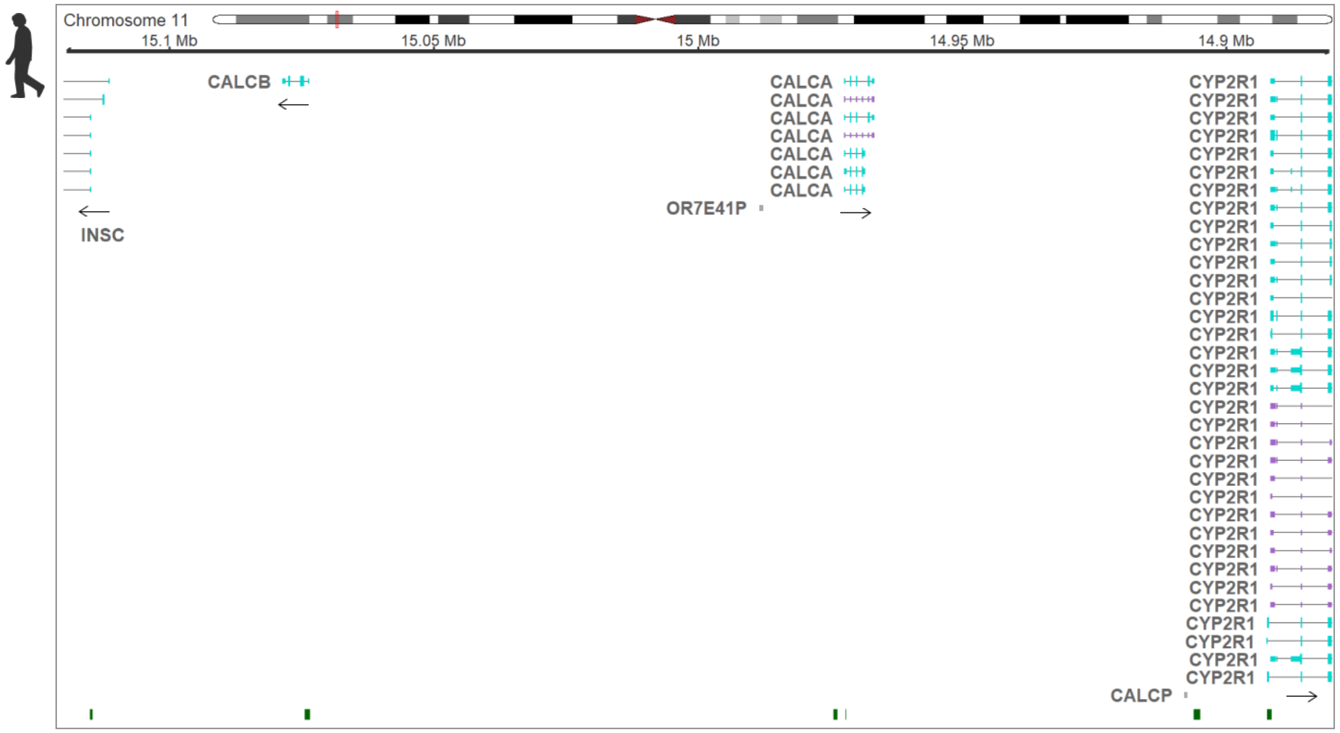


**Fig. S2. NCBI RefSeq gene model at the human *CALCB–CALCA* locus**. The full gene model between *INSC* and *CYP2R1* is shown, complementing Fig. 2B. *OR7341P* and *CALCP* are annotated as pseudogenes. CpG islands (CGIs) are displayed in the bottom track.

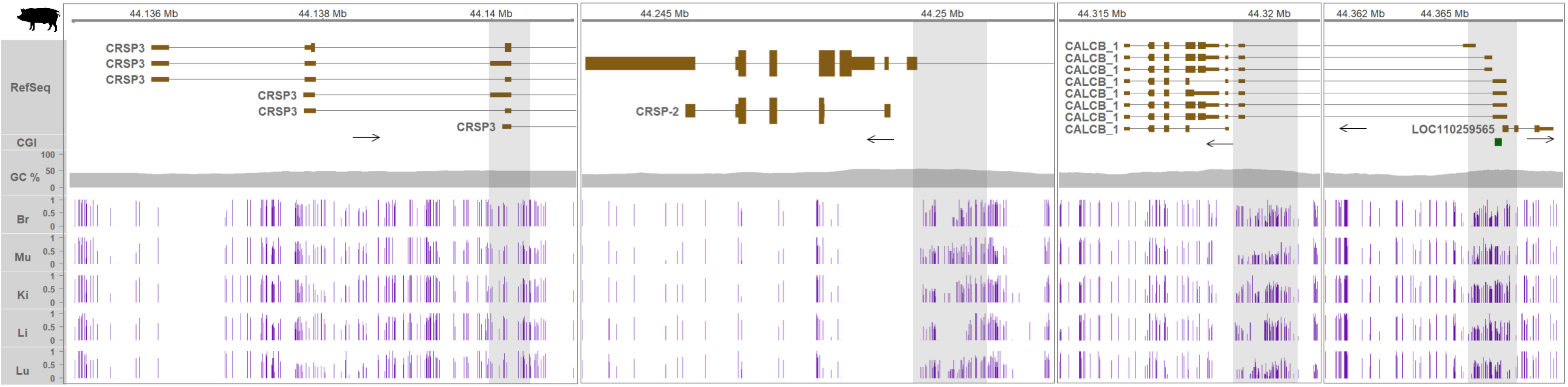


**Fig. S3. Zoomed views of DNA methylation in pig somatic tissues.** CpG methylation at putative promoter regions (grey shading) of CRSP-encoding genes, shown at a higher magnification than in Fig. 3. Transcriptional directions are indicated by arrows. Protein-coding transcripts are shown in brown. Tall and short boxes represent translated and untranslated regions, respectively. RefSeq, NCBI RefSeq gene model; CGI, CpG island; GC%, GC content; Br, brain; Mu, muscle; Ki, kidney; Li, liver; Lu, lung.

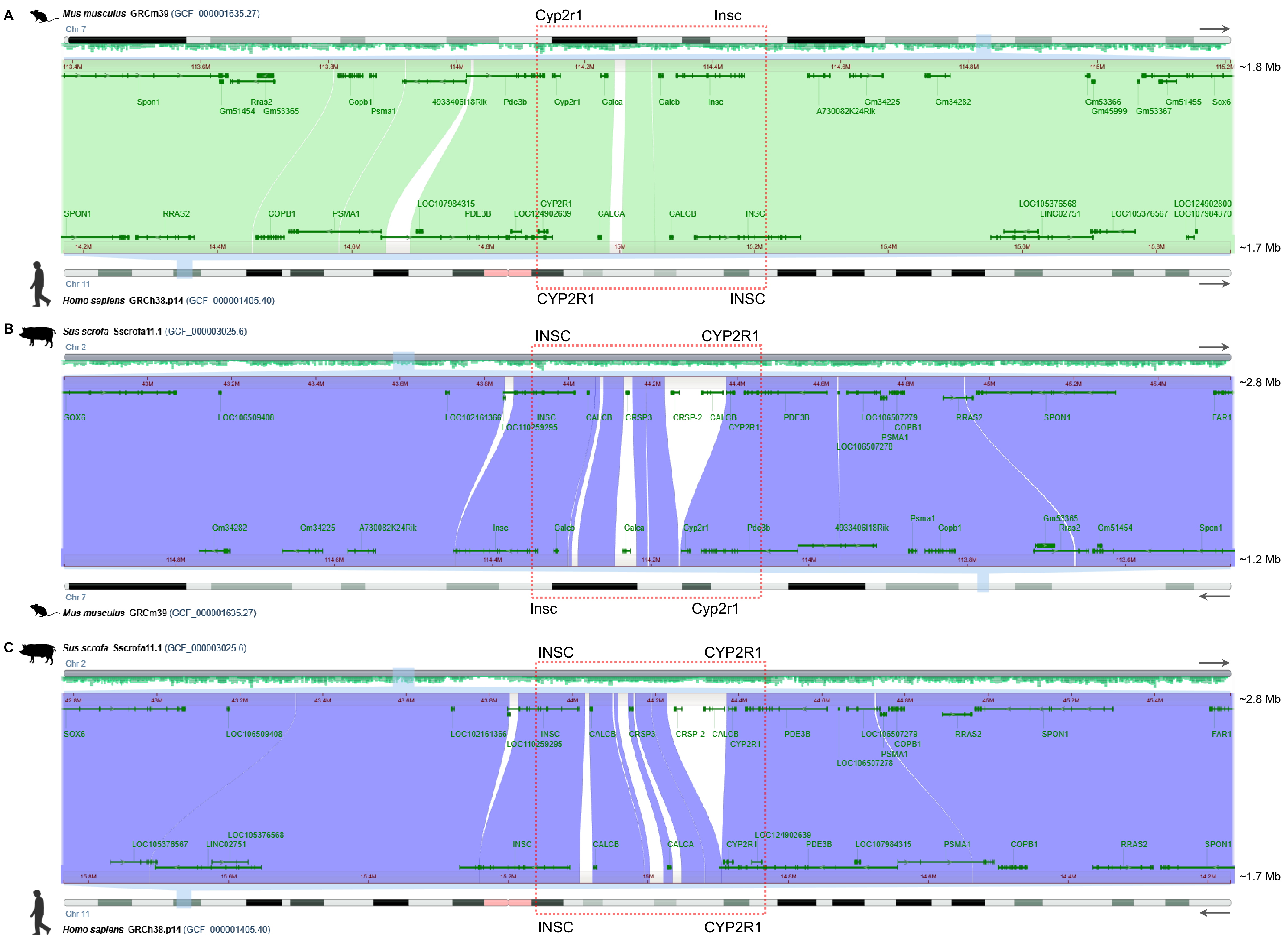


**Fig. S4. Synteny representation using the NCBI Comparative Genome Viewer (CGV).** Macrosyntenic views (approx. 1.2–2.8 Mb) of parts of mouse chromosome 7, human chromosome 11, and pig chromosome 2 are shown, highlighting the microsyntenic region encompassing *INSC* and *CYP2R1* (including the *CRSP* complex locus) with dashed boxes. Forward and reverse orientations of the chromosomes are indicated by arrows on the right. A comparison between mouse and human is shown with alignments in the forward orientation **(A)**, whereas comparisons between pig and mouse **(B)** and between pig and human **(C)** are displayed with alignments in the reverse orientation. Alignments in the forward and reverse orientations are visually distinguished by contrasting shading with purple in B and C.

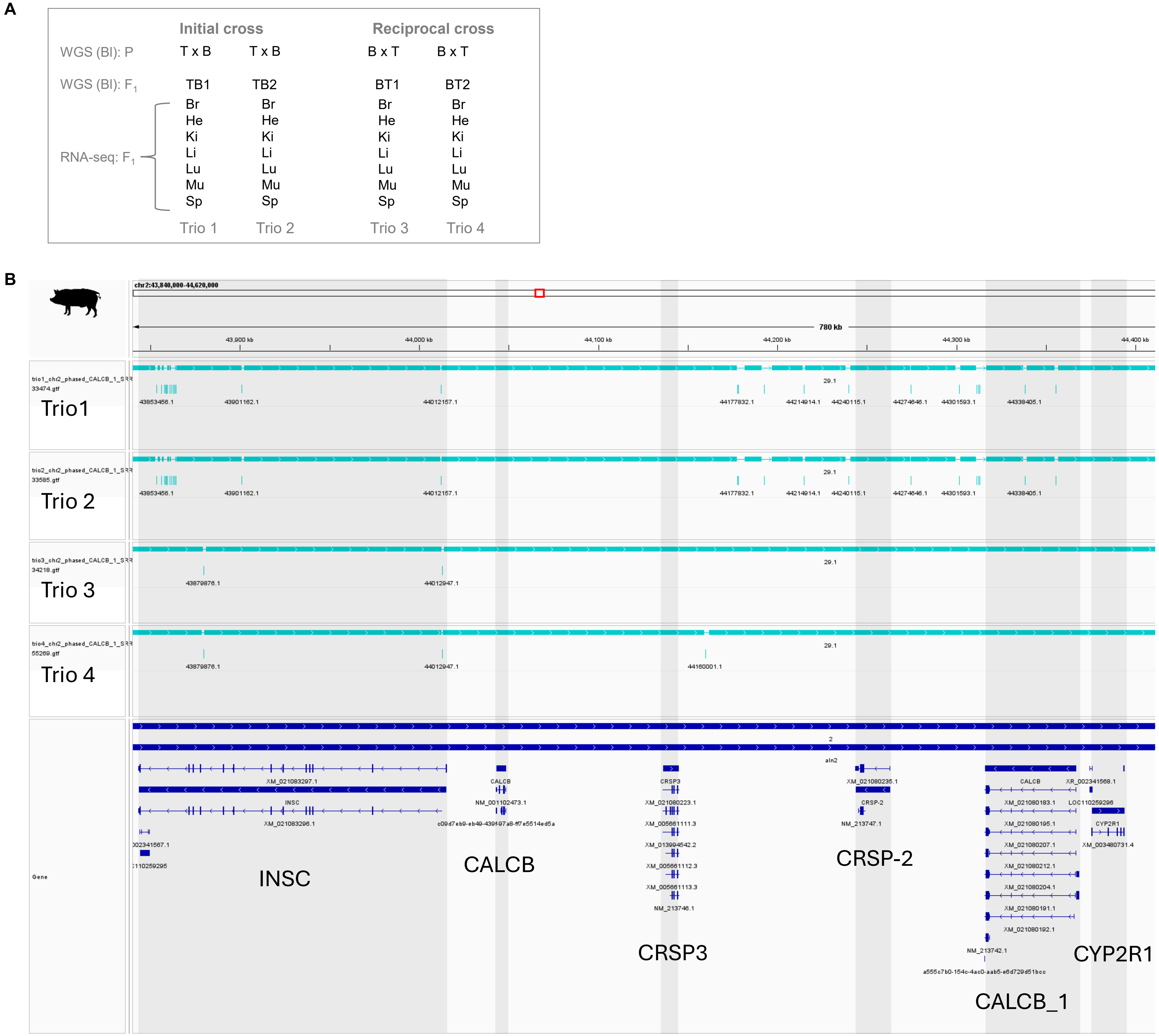


**Fig. S5. Cross design and phased blocks within the *INSC–CYP2R1* region. (A)** Structure of the data (initial and reciprocal crosses and trios; PRJNA998604) that were analyzed. T, Tibetan pigs; B, Berkshire pigs; TB, $F_1$ offspring of Tibetan sire × Berkshire dam cross (initial); BT, $F_1$ offspring of Berkshire sire × Tibetan dam cross (reciprocal); Bl, blood for WGS; Br, brain; He, heart; Ki, kidney; Li, liver; Lu, lung; Mu, muscle; Sp,spleen. **(B)** Phase sets (haplotype blocks; cyan bars) derived from WhatHap phase using WGS data.

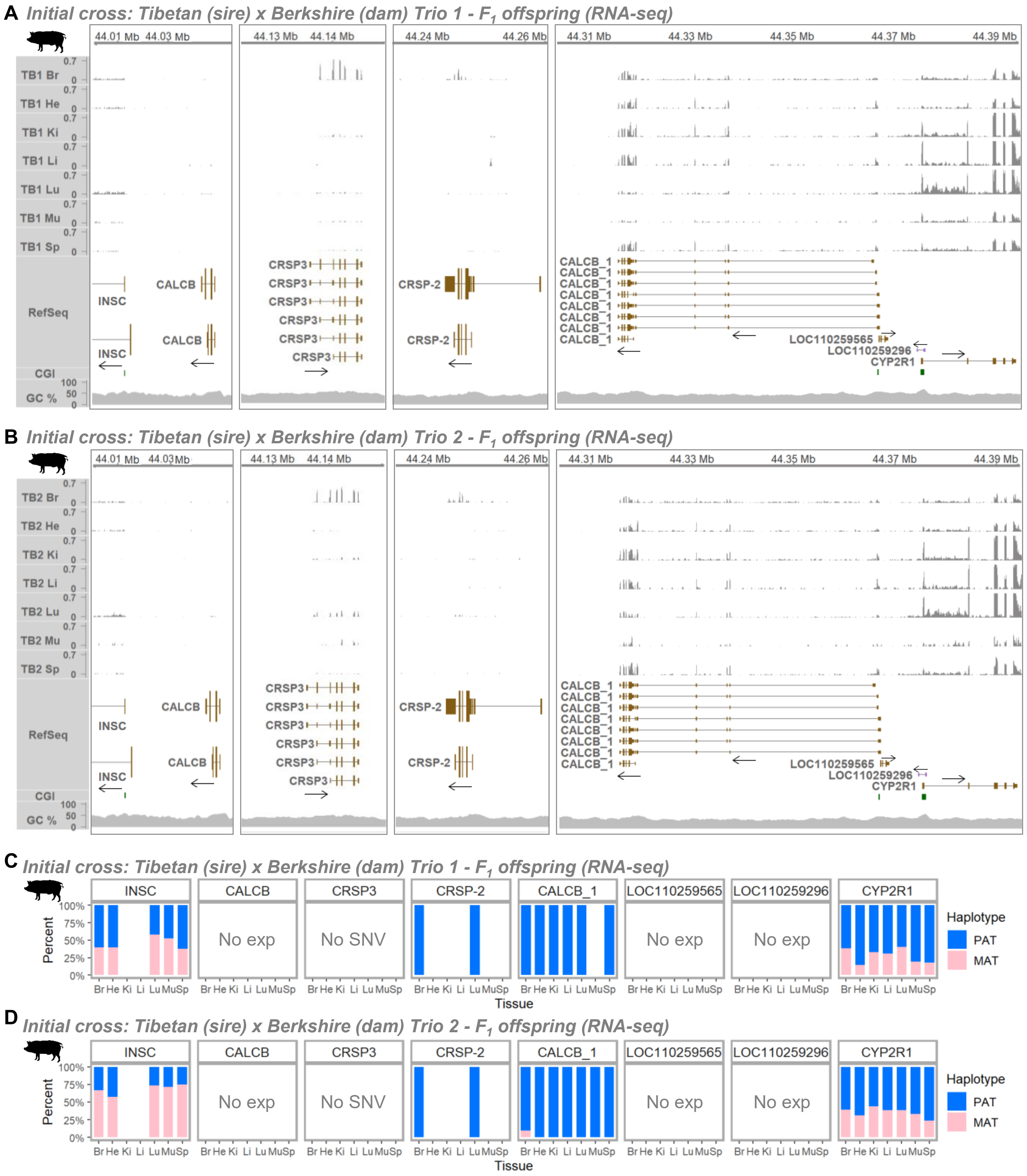


**Fig. S6. RNA expression of *CALCB*/*CRSP* genes and expressed parental haplotypes. (A, B)** RNA expression based on RNA-seq (PRJNA998604) in $F_1$ offspring from Trio 1 and 2 of the initial crosses. **(C, D)** Percentages of haplotype-tagged reads for paternal (PAT) and maternal (MAT) haplotypes, including only gene–tissue combinations with >5 haplotype-tagged reads (maximum depth: 379). T, Tibetan pigs; B, Berkshire pigs; Br, brain; He, heart; Ki, kidney; Li, liver; Lu, lung; Mu, muscle; Sp, spleen.

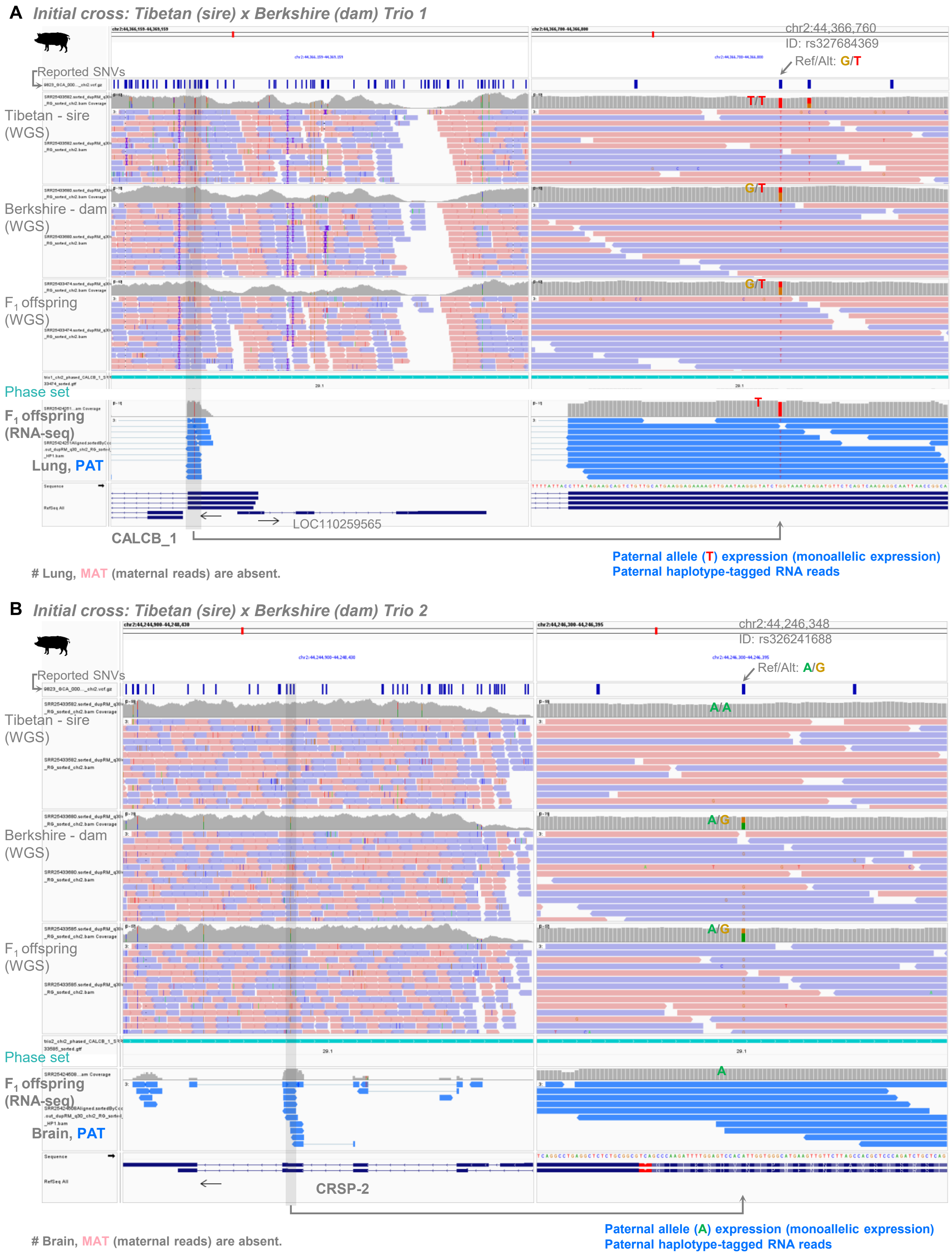


**Fig. S7. Exemplified IGV screenshots of haplotype phasing and tagging. (A)** *CALCB_1* in Trio 1 (initial cross). **(B)** *CRSP-2* in Trio 2 (initial cross). Reported SNVs were derived from the EBI EVA release 6 VCF for Sscrofa11.1.

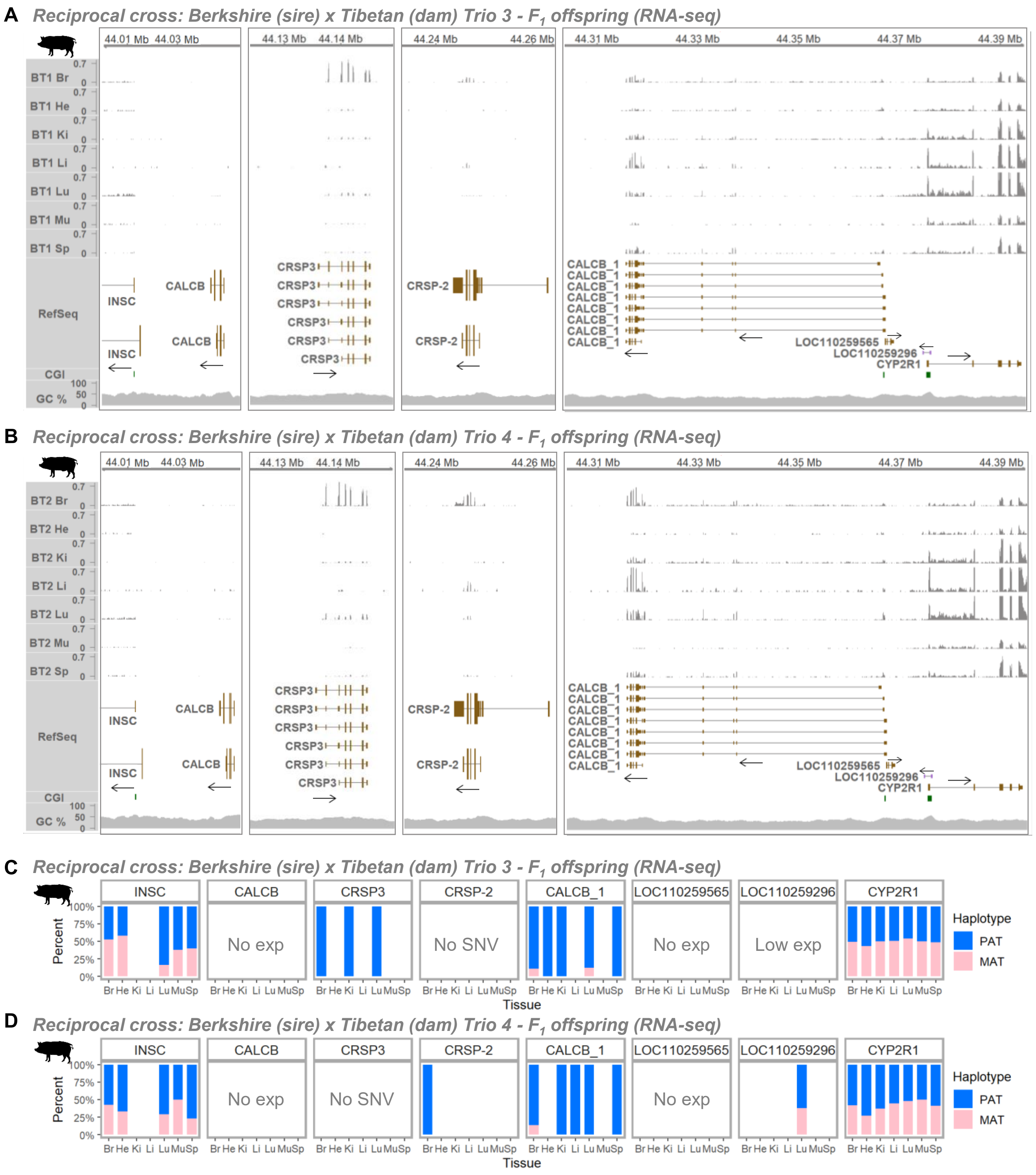


**Fig. S8. RNA expression of *CALCB*/*CRSP* genes and expressed parental haplotypes. (A, B)** RNA expression based on RNA-seq (PRJNA998604) in $F_1$ offspring from Trio 3 and 4 of the reciprocal crosses. **(C, D)** Percentages of haplotype-tagged reads for paternal (PAT) and maternal (MAT) haplotypes, including only gene–tissue combinations with >5 haplotype-tagged reads (maximum depth: 427). T, Tibetan pigs; B, Berkshire pigs; Br, brain; He, heart; Ki, kidney; Li, liver; Lu, lung; Mu, muscle; Sp, spleen.

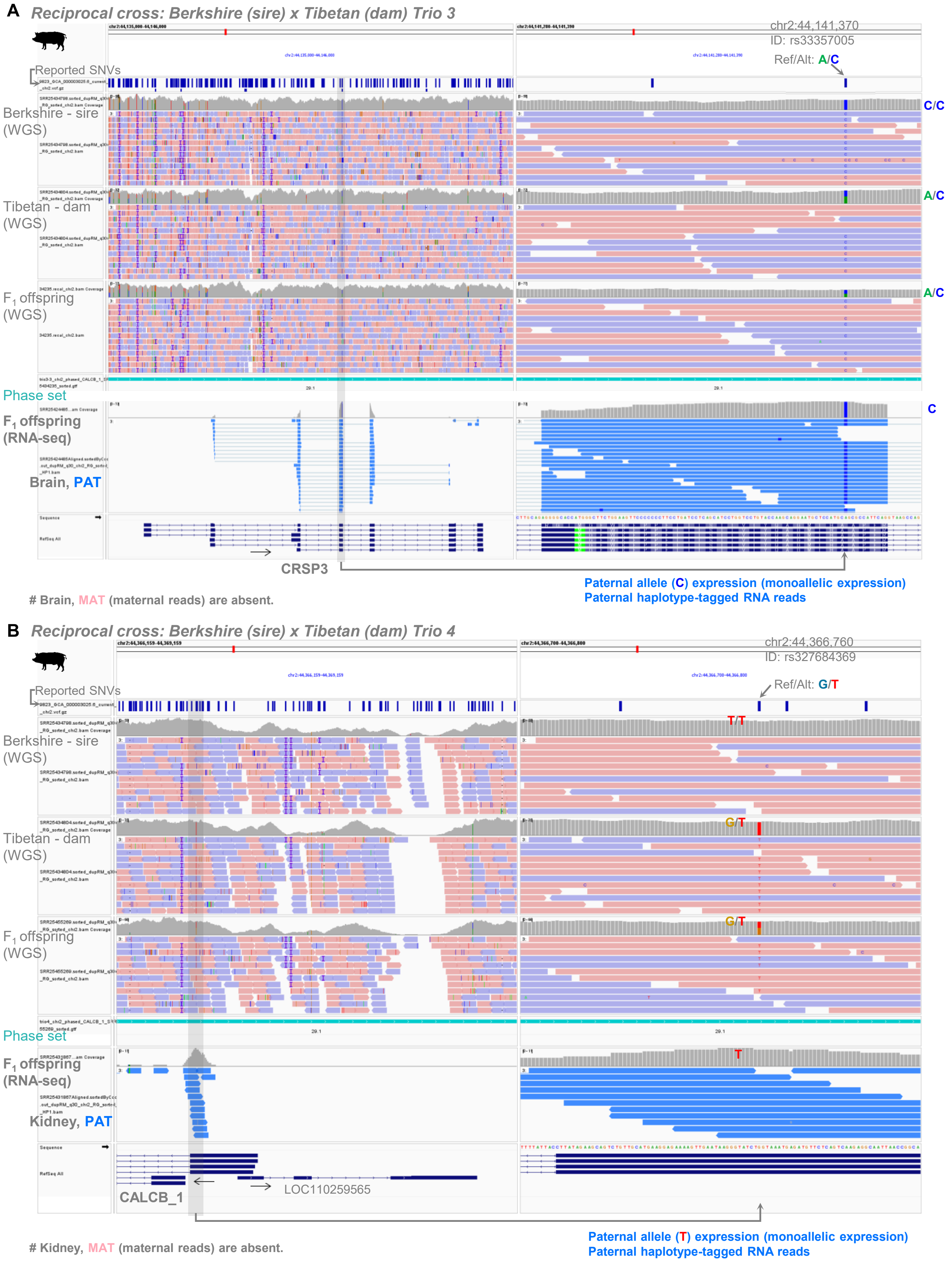


**Fig. S9. Exemplified IGV screenshots of haplotype phasing and tagging. (A)** *CRSP3* in Trio 3 (reciprocal cross). **(B)** *CALCB_1* in Trio 4 (reciprocal cross). Reported SNVs were derived from the EBI EVA release 6 VCF for Sscrofa11.1.

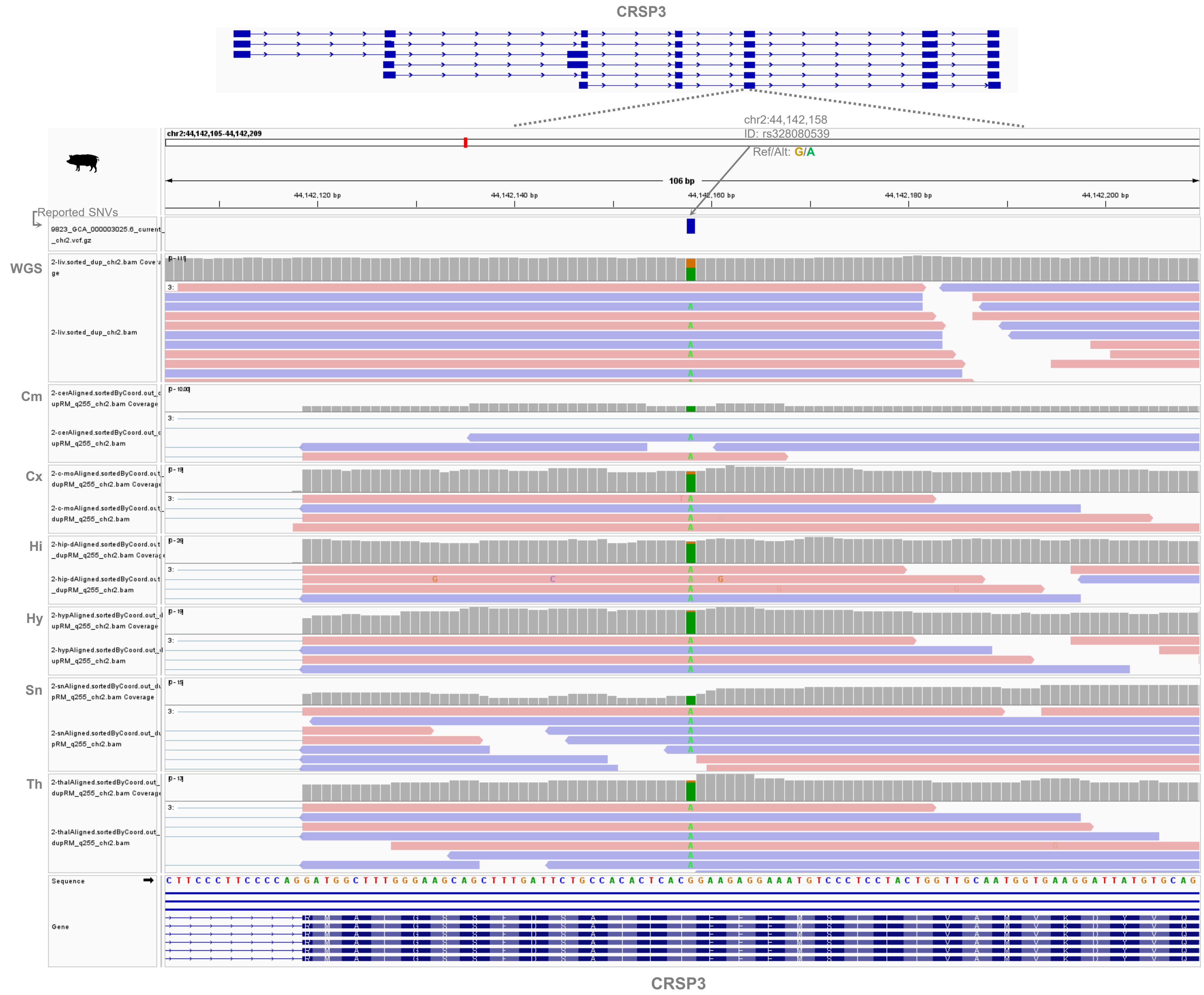


**Fig. S10. Allelic expression in brain subregions.** A heterozygous SNV located in an exon of *CRSP3* and the expressed allele in brain subregions are shown. WGS and RNA-seq data were derived from the same 1-year-old Bama minipig. Cm, cerebellum; Cx, motor cortex, Hi, hippocampus, Hy, hypothalamus, Sn, substantia nigra; Th, thalamus. The WGS (CNP0001045) and RNA-seq (CNP0000483) datasets were retrieved from the CNGBdb repository. Reported SNVs were derived from the EBI EVA release 6 VCF for Sscrofa11.1.

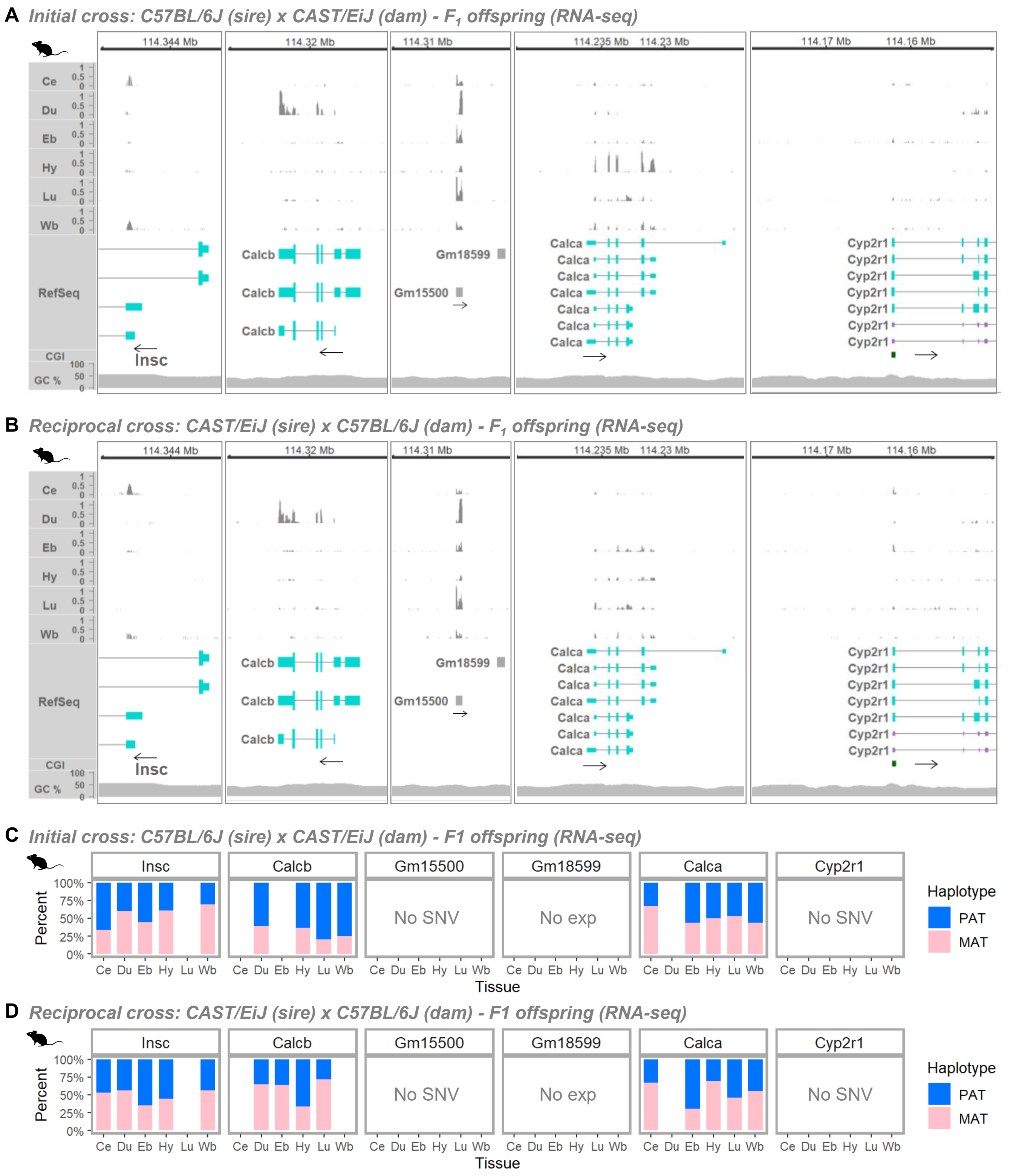


**Fig. S11. RNA expression of *Calcb* and *Calca* and expressed parental haplotypes. (A, B)** RNA expression based on RNA-seq in $F_1$ offspring at embryonic or adult stages from initial and reciprocal crosses (accession SRP020526). **(C, D)** Percentages of haplotype-tagged reads for paternal (PAT) and maternal (MAT) haplotypes, including only gene–tissue combinations with >5 haplotype-tagged reads (maximum depth: 136). Ce, cerebellum; Du, duodenum; Eb, embryonic brain; Hy, hypothalamus; Lu, lung; Wb, whole brain. No SNV, absence of SNVs in CAST/EiJ mice; No exp, no detectable RNA expression across tissues.

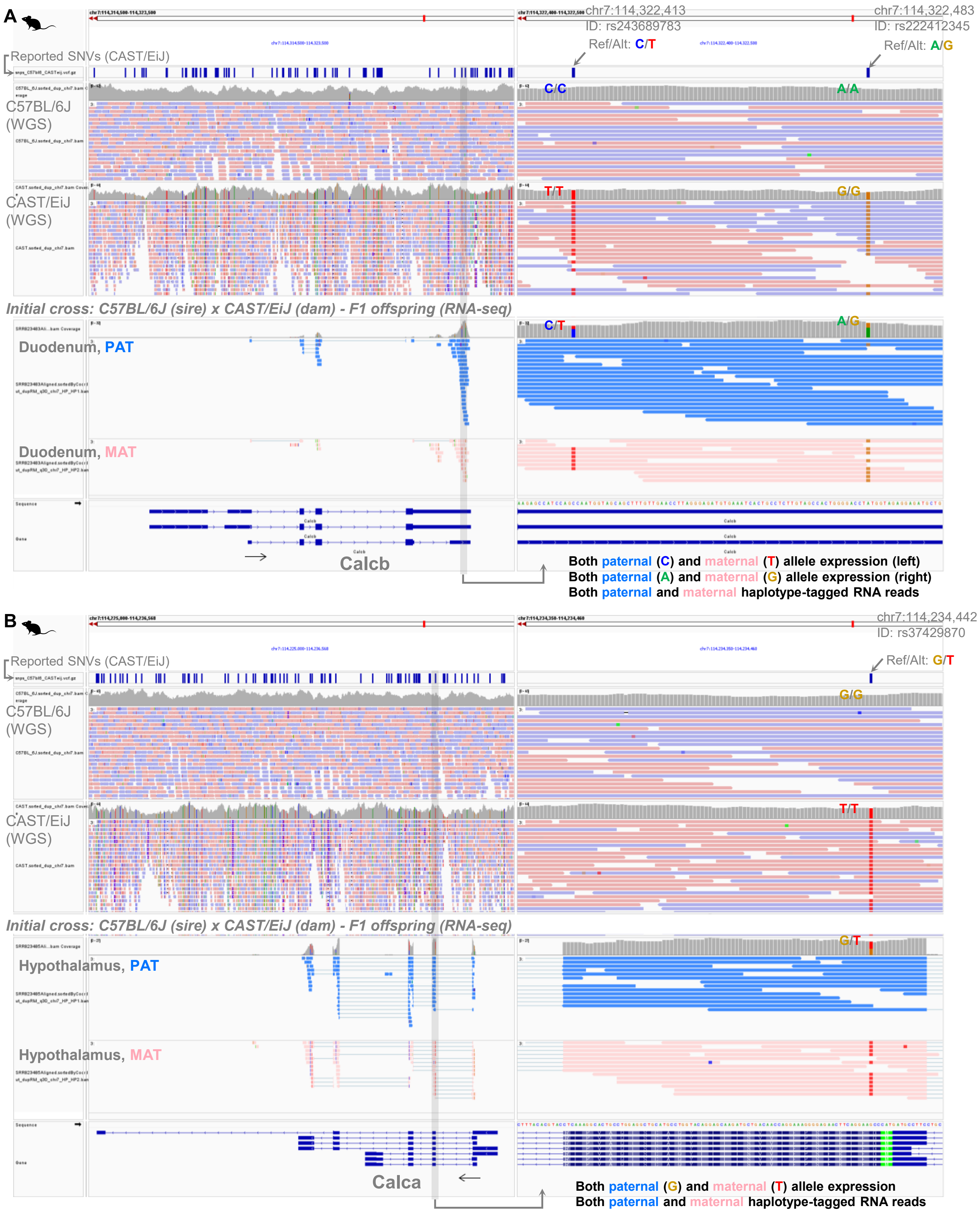


**Fig. S12. Exemplified IGV screenshots of haplotype tagging in the initial cross. (A)** *Calcb* in the initial cross. **(B)** *Calca* in the initial cross. The WGS tracks for C57BL/6J and CAST/EiJ are shown only to illustrate strain-specific genomic sequences and were not used for phasing. Instead, haplotype tagging of RNA-seq reads was performed using the pseudo-phased VCF generated from strain-specific variants, as described in the Results section. Reported SNVs were derived from the Mouse Genomes Project (MGP; REL2021).

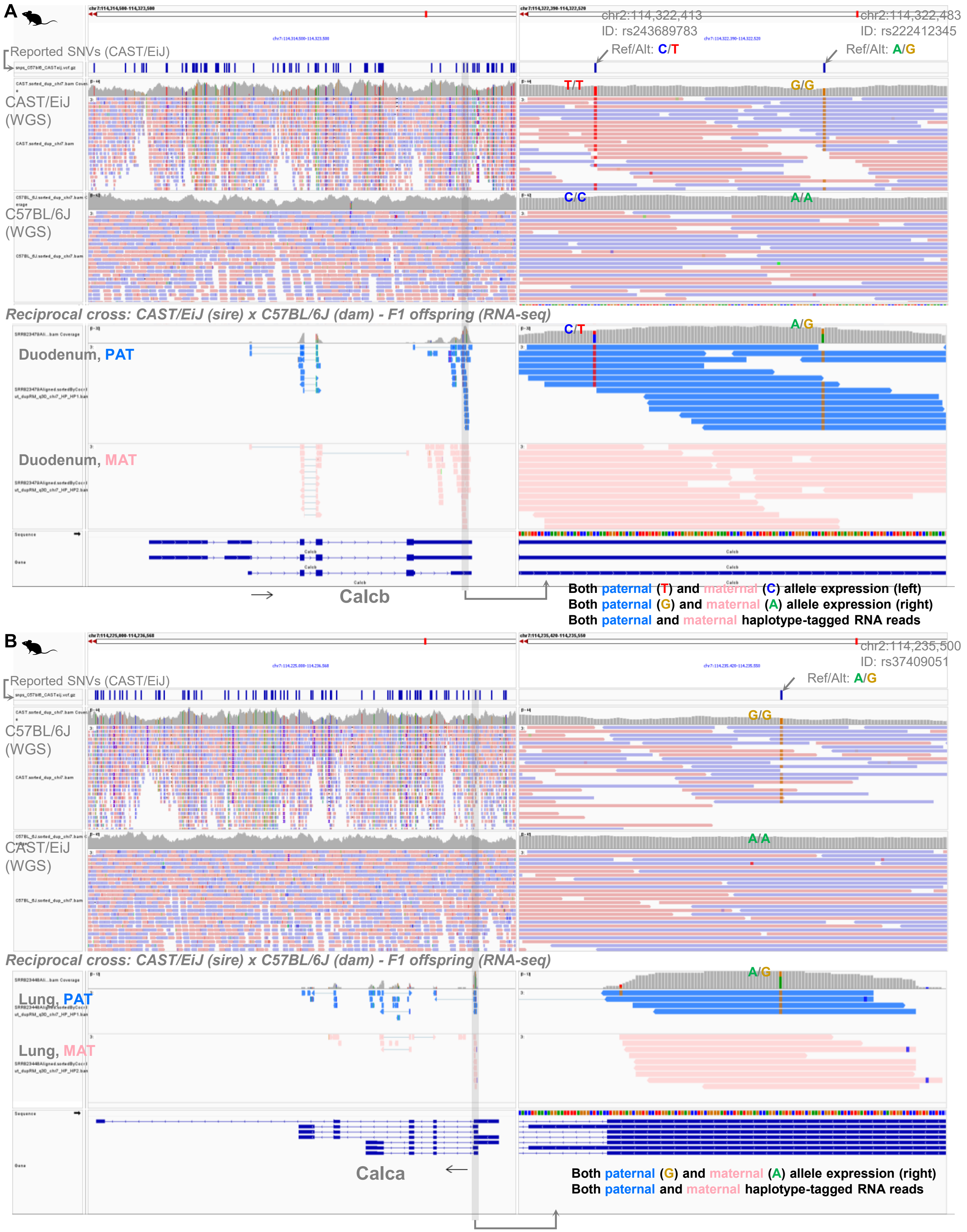


**Fig. S13. Exemplified IGV screenshots of haplotype tagging in the reciprocal cross. (A)** *Calcb* in the reciprocal cross. **(B)** *Calca* in the reciprocal cross. The WGS tracks for CAST/EiJ and C57BL/6J are shown only to illustrate strain-specific genomic sequences and were not used for phasing. Instead, haplotype tagging of RNA-seq reads was performed using the pseudo-phased VCF generated from strain-specific variants, as described in the Results section. Reported SNVs were derived from the Mouse Genomes Project (MGP; REL2021).

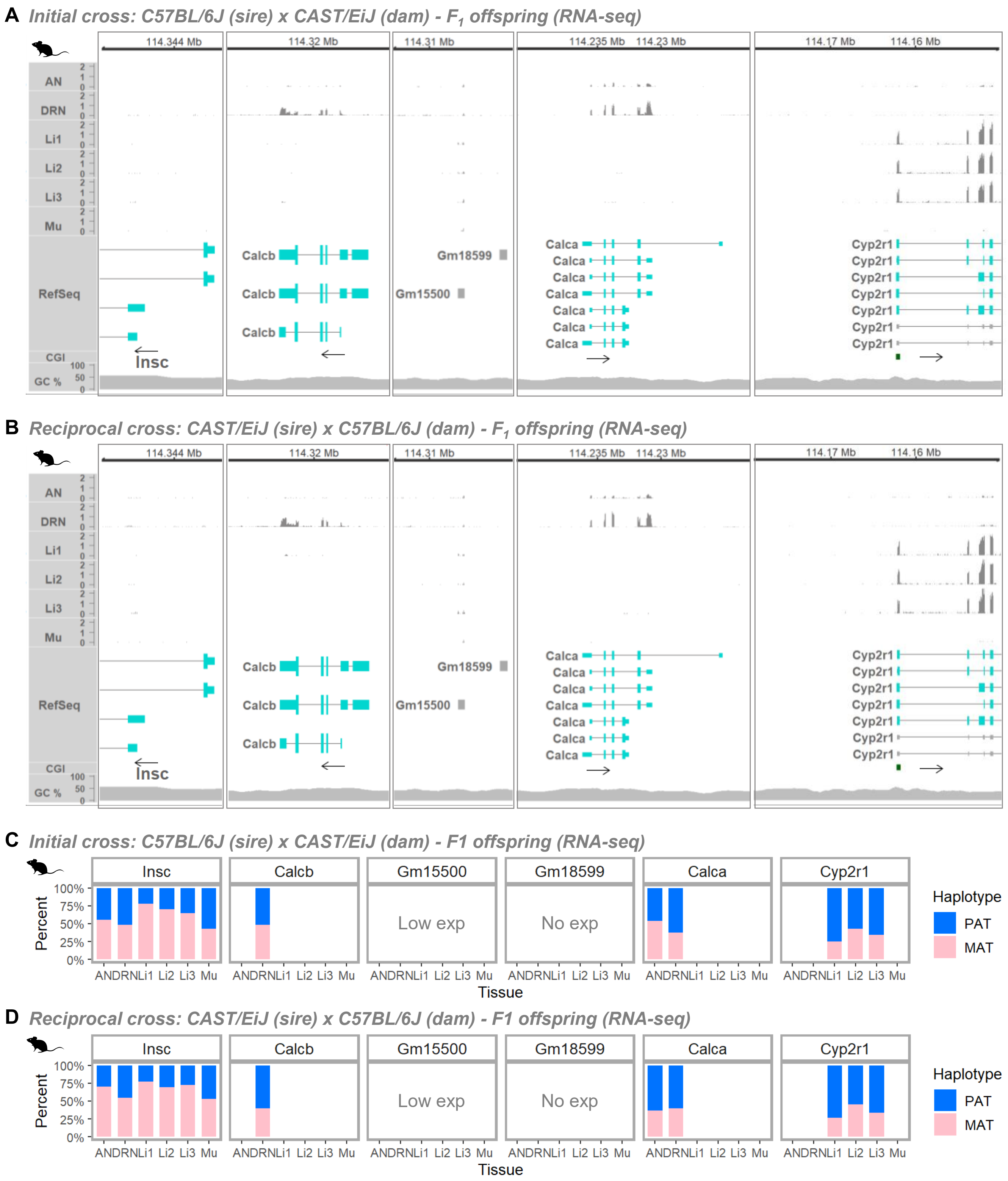


**Fig. S14. RNA expression of *Calcb*, *Calca*, and *Cyp2r1*, and expressed parental haplotypes. (A, B)** RNA expression based on RNA-seq in $F_1$ offspring (8 weeks old) from initial and reciprocal crosses (accession GSE70484). **(C, D)** Percentages of haplotype-tagged reads for paternal (PAT) and maternal (MAT) haplotypes, including only gene–tissue combinations with >5 haplotype-tagged reads (maximum depth: 486). AN, arcuate nucleus; DRN, dorsal raphe nucleus; Li, liver; Mu, muscle. Low or No exp, low or no detectable RNA expression across tissues.

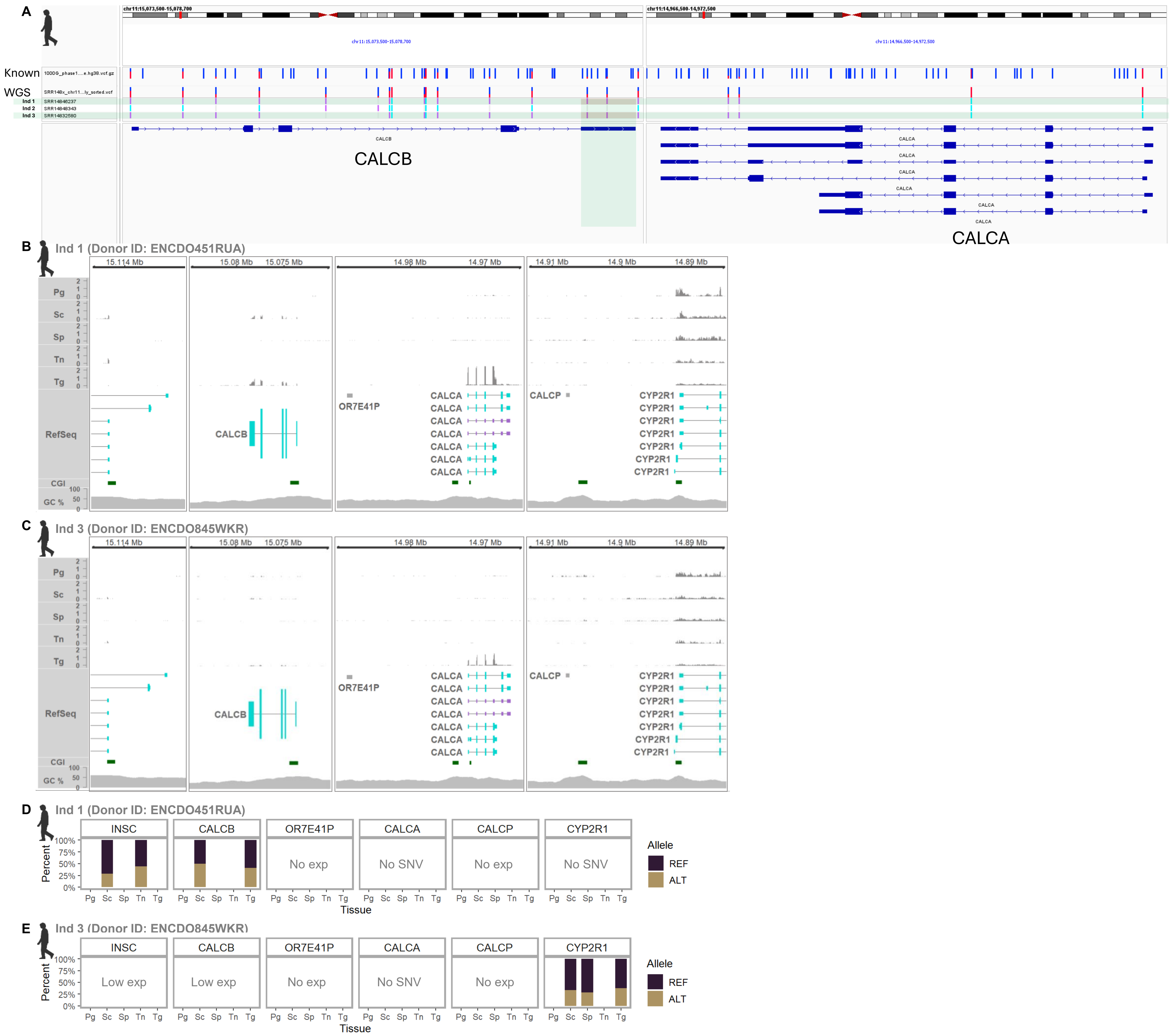
A
Known
WGS
Ind 1
Ind 2
Ind 3
CALCB
CALCA
B
Ind 1 (Donor ID: ENCDO451RUA)
Pg
Sc
Sp
Tn
Tg
RefSeq
CGI
GC %
OR7E41P
CALCP
CYP2R1
C
Ind 3 (Donor ID: ENCDO845WKR)
D
Ind 1 (Donor ID: ENCDO451RUA)
INSC
CALCB
OR7E41P
CALCA
CALCP
CYP2R1
No exp
No SNV
Percent
Tissue
Allele
REF
ALT
E
Ind 3 (Donor ID: ENCDO845WKR)
Low exp

F

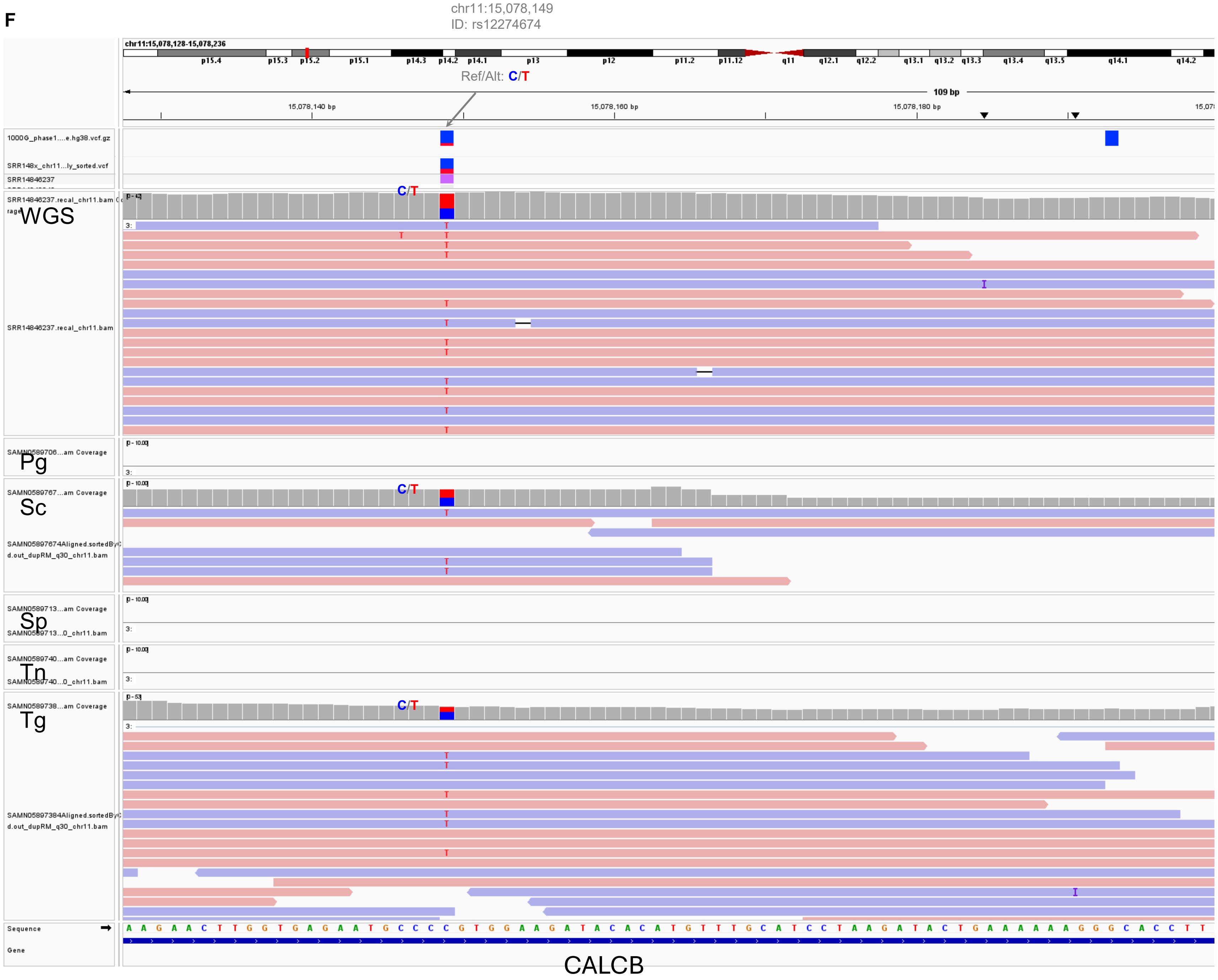


**Fig. S15. SNV calling and analysis of allelic expression using human ENCODE data. (A)** WGS data from three human individuals ("Ind") were analyzed (PRJNA63443; donor IDs: ENCDO451RUA (54 years, top), ENCDO793LXB (53 years, middle), and ENCDO845WKR (37 years, bottom)). Heterozygous SNVs (purple bars) on the last exon of *CALCB* are highlighted with green and red shading in Ind 1 (ENCDO451RUA) and Ind 3 (ENCDO845WKR), and homozygous SNVs are denoted with cyan bars. Known variant sites ("Known") were retrieved from 1000G_phase1.snps.high_confidence.hg38.vcf.gz (Broad Institute hg38 resource bundle). **(B-C)** Corresponding RNA-seq from ENCDO451RUA and ENCDO845WKR. Pg, prostate gland; Sc, sigmoid colon; Sp, spleen; Tn, tibial nerve; Tg, thyroid gland. **(D-E)** Percentages of allele counts for ENCDO451RUA and ENCDO845WKR. REF, reference allele; ALT, alternative allele. **(F)** Representative IGV screenshot for rs12274674 (chr11:15,078,149) in ENCDO451RUA (Ind 1).

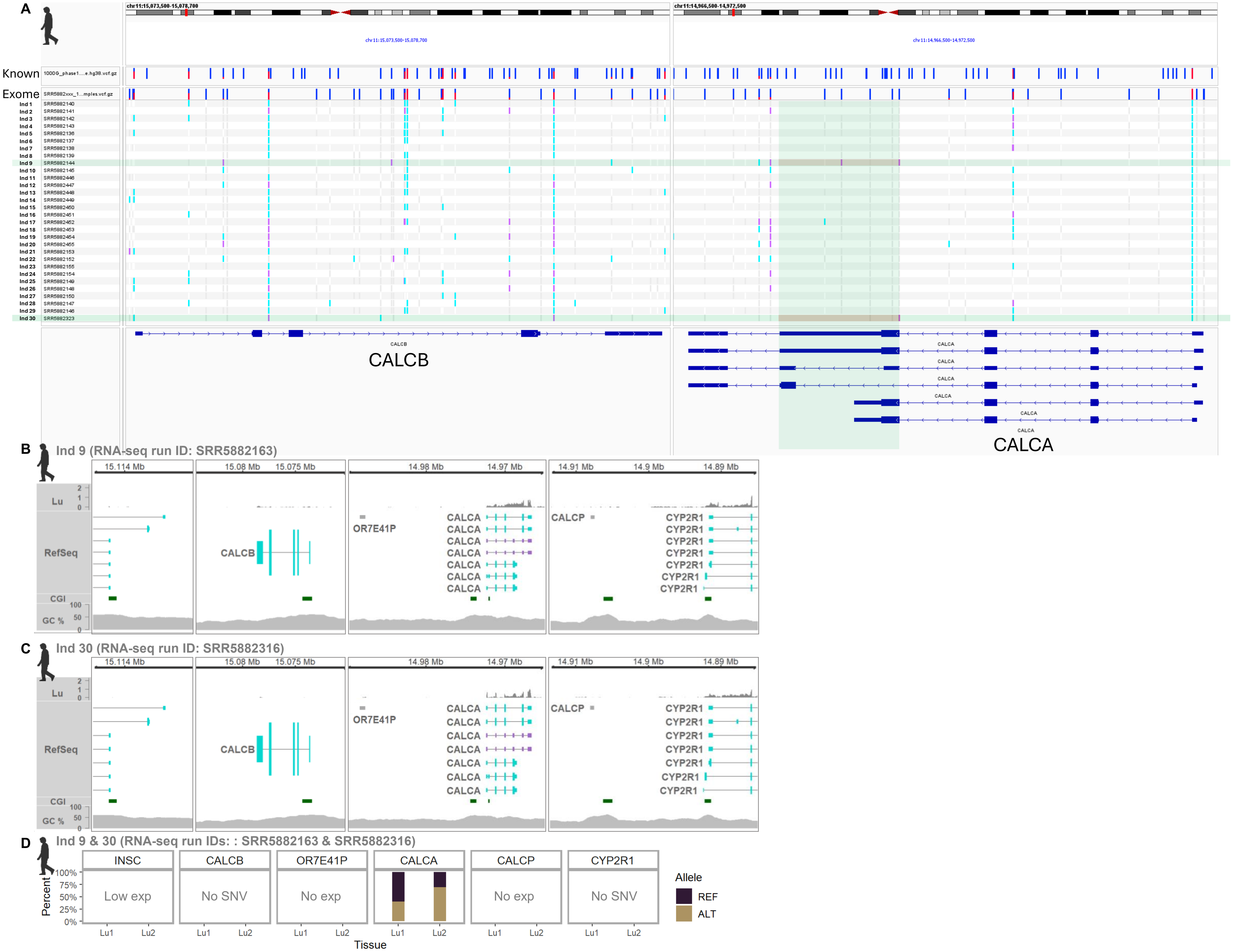

A
Known
Exome
chr11:15,073,500-15,078,700
chr11:14,966,500-14,972,500
CALCB
CALCA
B
Ind 9 (RNA-seq run ID: SRR5882163)
C
Ind 30 (RNA-seq run ID: SRR5882316)
Lu
RefSeq
CGI
GC %
15.114 Mb
15.08 Mb
15.075 Mb
14.98 Mb
14.97 Mb
14.91 Mb
14.9 Mb
14.89 Mb
OR7E41P
CALCP
CYP2R1
D
Ind 9 & 30 (RNA-seq run IDs: : SRR5882163 & SRR5882316)
INSC
Low exp
No SNV
No exp
Percent
Lu1
Lu2
Tissue
Allele
REF
ALT

E

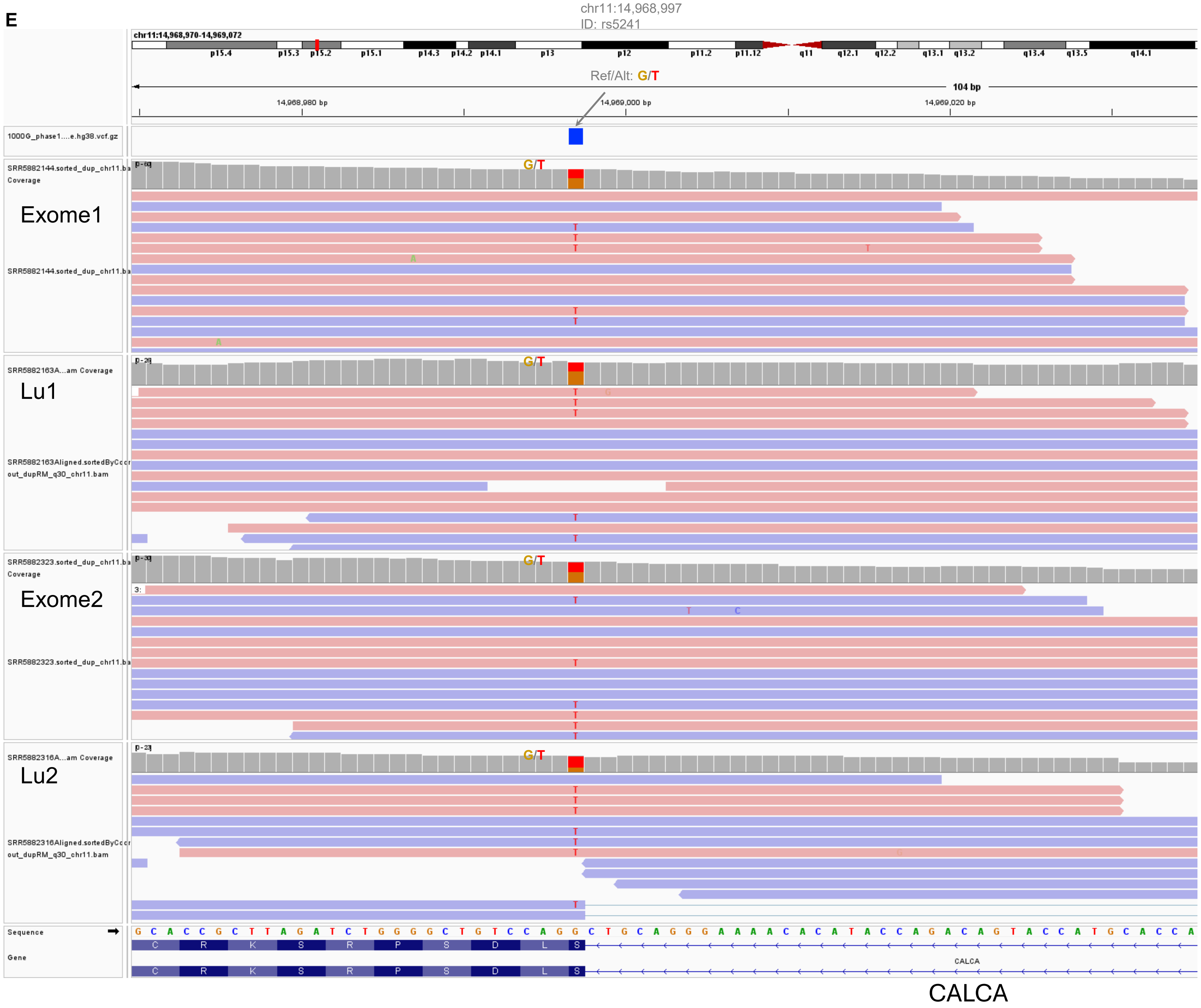


**Fig. S16. SNV calling and analysis of allelic expression using human lung data. (A)** Exome data from 30 human individuals ("Ind") were analyzed (PRJNA395106; exome run IDs: SRR5882136–SRR5882150, SRR5882152–SRR5882155, SRR5882323, SRR5882446–SRR5882455; age range: 35–83). Heterozygous SNVs (purple bars) on the exon are highlighted with green and red shading in Ind 9 (SRR5882144) and Ind 30 (SRR5882323), and homozygous SNVs are indicated with cyan bars. Known variant sites ("Known") were obtained from 1000G_phase1.snps.high_confidence.hg38.vcf.gz (Broad Institute hg38 resource bundle). **(B-C)** Corresponding RNA-seq from Ind 9 (SRR5882163) and Ind 30 (SRR5882316). Lu, lung. **(D)** Allele counts for Ind 9 and Ind 30. REF, reference allele; ALT, alternative allele. **(E)** Representative IGV screenshot for rs5241 (chr11:14,968,997) in Ind 9 and Ind 30.

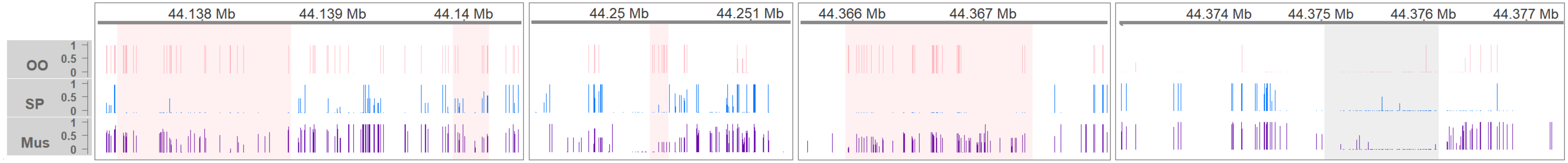


**Fig. S17. Zoom-in view of the regions highlighted in the WGBS data presented in Fig.6D.** The regions marked with red bars in Fig. 6D corresponding to *CRSP3*, *CRSP-2*, and *CALCB_1* are enlarged in the left three panels. The rightmost panel shows the first exon of *CYP2R1*, highlighted with grey shading.

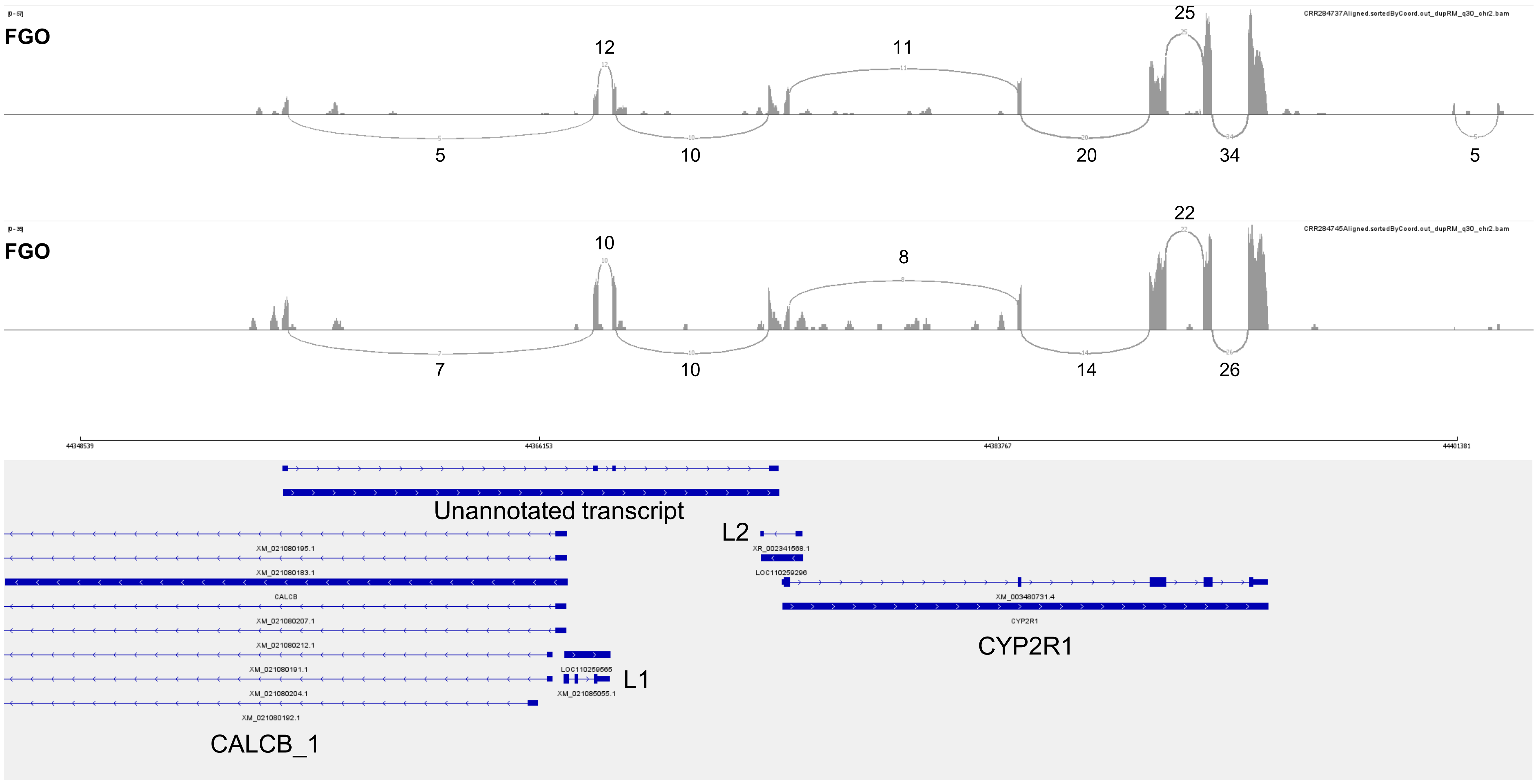


**Fig. S18. Sashimi plot for the unannotated transcript between *CALCB_1* and *CYP2R1* in pig oocytes.** The IGV-based sashimi plot of RNA-seq reads (GSE163709) is presented with numerical read coverage for each splicing junction. The annotated *CYP2R1* transcript is also shown in the downstream region.

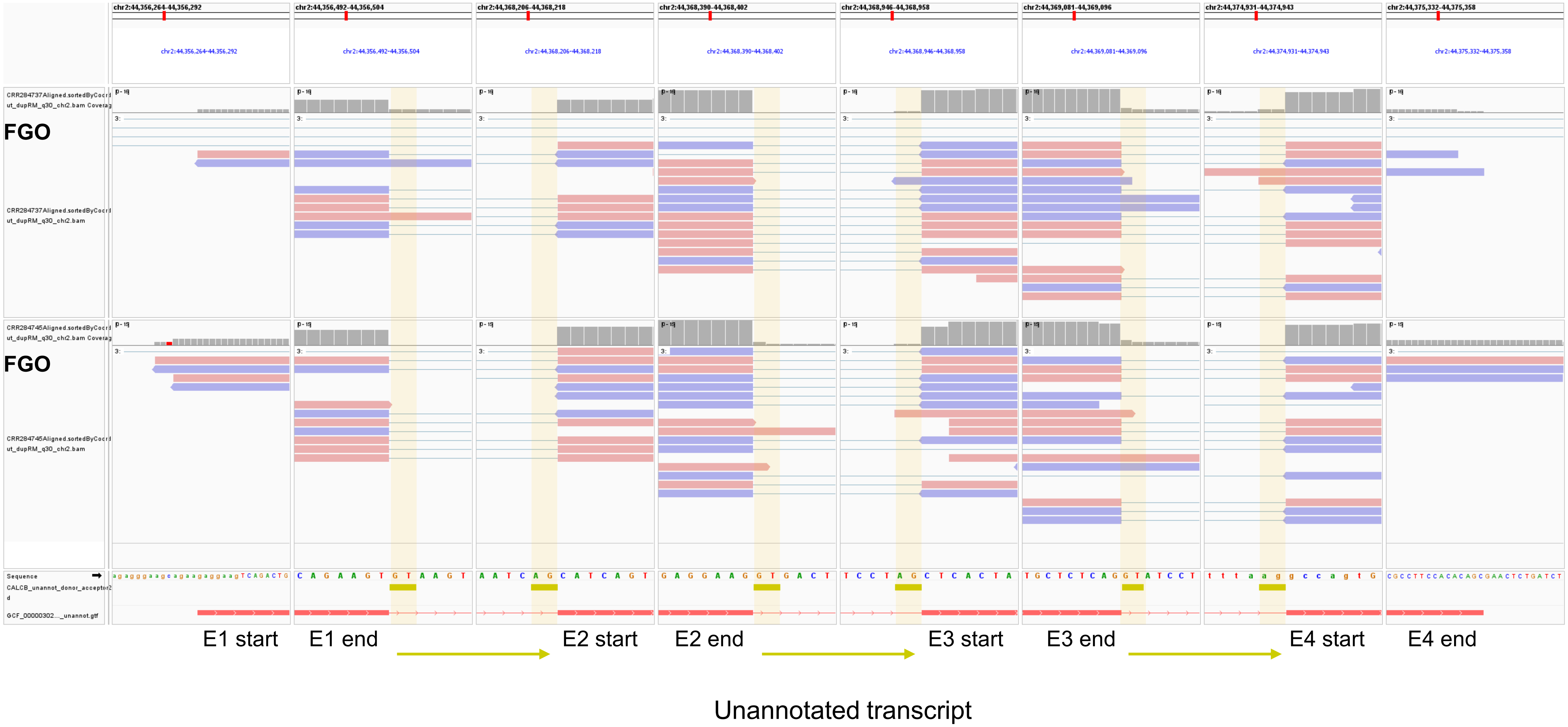


**Fig. S19. Predicted transcriptional direction of the unannotated transcript between *CALCB_1* and *CYP2R1* in pig oocytes.** The genomic positions of splice donor (GT) and splice acceptor (AG) sites are indicated with yellow bars and shading.
